# Characterization of *Drosophila Nidogen/entactin* reveals roles in basement membrane stability, barrier function and nervous system plasticity

**DOI:** 10.1101/348631

**Authors:** Georg Wolfstetter, Ina Dahlitz, Kathrin Pfeifer, Joscha Arne Alt, Uwe Töpfer, Daniel Christoph Pfeifer, Reinhard Lakes-Harlan, Stefan Baumgartner, Ruth H. Palmer, Anne Holz

**Author notes:** Both authors contributed equally (Ina Dahlitz, Kathrin Pfeifer).

## Abstract

Basement membranes (BMs) are specialized layers of extracellular matrix (ECM) mainly composed of Laminin, type IV Collagen, Perlecan and Nidogen/entactin (NDG). While the essential and evolutionary conserved functions of Laminin, Collagen and Perlecan are well documented in *Drosophila* and other species, the proposed role of NDG as the major ECM linker molecule has been challenged by several *in vivo* studies revealing that NDG is dispensable for viability and BM formation. Here, we report the characterization of the single *Ndg* gene in *Drosophila.* Embryonic *Ndg* expression differed from that of other BM components and was primarily observed in mesodermal tissues and the chordotonal organs, whereas NDG protein localized to all BMs. While loss of Laminin strongly affected BM-localization of NDG, *Ndg* null mutants exhibited no overt changes in the distribution of BM core components. However, loss of NDG led to ultrastructural BM defects compromising barrier function and stability *in vivo.* Although *Ndg* mutants were viable, loss of NDG led to decreased fecundity in flies as well as impaired crawling behavior and reduced response to vibrational stimuli in larvae. Further morphological analysis revealed accompanying defects in the larval peripheral nervous system especially in the chordotonal organs and the neuromuscular junction (NMJ), where *Ndg* genetically interacted with the *Leukocyte-antigen-related-like (Lar) receptor* gene to regulate NMJ extension and synaptic differentiation. Taken together, our analysis suggests that NDG is not essential for BM assembly but mediates BM stability and ECM-dependent neural plasticity during *Drosophila* development.

**Summary Statement:** In this study we characterize *Drosophila Nidogen/Entactin (Ndg)* mutants revealing that loss of Ndg impairs basement membrane (BM) stability and permeability as well as proper function of the nervous system.

## Introduction

Proteome analysis of isolated BMs has identified over 100 associated proteins highlighting the complex nature of these specialized ECM sheets (Uechi et al., 2014). However, BMs mainly assemble from a small subset of ECM proteins referred to as the “BM toolkit” which is a highly conserved feature of most metazoan species (Hynes, 2012). BMs consist of two different mesh-like networks formed by self-assembly of either heterotrimeric Laminin molecules or type IV Collagens which are then linked by their binding partners Nidogen (NDG), the heparan sulfate proteoglycan (HSPG) Perlecan and additional Collagen XV/XVIII homologs (Jayadev and Sherwood, 2017; Yurchenco, 2011). The *Drosophila* genome harbors a minimal set of nine genes encoding components of the “BM toolkit” (Hynes, 2012). Four genes encode for two Laminin α-, one β- and one γ-subunit which form the only two Laminin heterotrimers in the fly, two adjacent loci encode for type IV Collagens (*Col4a1* and *viking*), one for a ColXV/XVIII homolog named *Multiplexin*, a single *Nidogen/entactin* gene, and the *terribly reduced optic lobes* (*trol*) locus that encodes for the Perlecan protein core. With the exception of *Ndg*, all mayor BM components of *Drosophila* have been functionally characterized (Borchiellini et al., 1996; Datta and Kankel, 1992; Friedrich et al., 2000; Henchcliffe et al., 1993; Lindsley and Zimm, 1992; Martin et al., 1999; Meyer and Moussian, 2009; Urbano et al., 2009; Voigt et al., 2002; Wolfstetter and Holz, 2012; Yarnitzky and Volk, 1995; Yasothornsrikul et al., 1997).

In 1983, Timpl et al (Timpl et al., 1983) reported the identification of an 80 kDA protein from mouse Engelbreth-Holm-Swarm (EHS) sarcoma cell lines which was named Nidogen due to its ability to self-aggregate into “nest-like structures”. Although this protein was later identified as a proteolytic fragment of Entactin, a 150 kDa sulfated glycoprotein described by Carlin et al. (Carlin et al., 1981), the terms Nidogen or Nidogen/Entactin are now commonly used to refer to the non-cleaved protein (Martin and Timpl, 1987). The Nidogen protein consists of three globular domains (G1 to G3), a rod-like segment between G1 and G2 as well as an EGF-like-domain-containing segment that connects G2 and G3 (Durkin et al., 1988; Fox et al., 1991). Nidogen forms stable complexes with Laminin through binding of its globular G3 domain to the Laminin γ-subunit, whereas the G2 domain mediates binding to Collagen IV and Perlecan (Fox et al., 1991; Hopf et al., 2001; Mann et al., 1989; Reinhardt et al., 1993). Therefore it has been suggested that Nidogen supports the formation of ternary complexes within the BM and functions as an “ECM linker molecule” connecting the Laminin and Collagen IV networks (Aumailley et al., 1993; Ho et al., 2008; Mayer et al., 1993; Mayer et al., 1995).

Functional analysis of the two Nidogen family members in mammals reveals that the single knockouts of *Nidogen 1* (*NID1*) and *Nidogen 2* (*NID2*) in mice neither result in lethality nor cause any gross defects in BM formation and morphology. However, *NID1^−/−^.* animals exhibit neurological phenotypes like spontaneous seizures and hind limb ataxia (Dong et al., 2002; Murshed et al., 2000; Schymeinsky et al., 2002). Moreover, elevated NID2 levels in BMs of *NID1^−/−^.* mice suggest a functional redundancy between the two members of the mammalian Nidogen family (Miosge et al., 2000). Analysis of double mutant *NID1^−/−^. NID2^−/−^.* mice does not reveal an essential function during embryogenesis and embryonic BM formation but complete absence of Nidogen causes perinatal lethality due to impaired lung and heart development accompanied by defects in the organ-associated BMs (Bader et al., 2005). Moreover, a varying degree of syndactyly and occasional twisting of forelimbs is observed in *NID1^−/−^. NID2^−/−^.* animals (Bose et al., 2006). In accordance with these results a non-essential function of Nidogen for invertebrate development is revealed by the analysis of *C. elegans nidogen-1 (nid-1*, the single *Nidogen* homolog in the nematode) mutants that are viable and fertile and do not display abnormalities during BM assembly. In addition to a slight reduction in fecundity, *nid-1.* mutant animals exhibit aberrant guidance and positioning of longitudinal nerves, movement defects in a body bending assay, and altered neuromuscular junction (NMJ) organization which suggests a role in nervous system patterning (Ackley et al., 2003; Hobert and Bulow, 2003; Kang and Kramer, 2000; Kim and Wadsworth, 2000; Kramer, 2005). A recent study by Zu et al. (Zhu et al., 2017) reports body length reduction in *Danio rerio* upon *nid1a* gene depletion but also suggests that Ndg-associated phenotypes are likely to be obscured by functional redundancy and genetic compensation (Rossi et al., 2015) between the four predicted *Nidogen* family members in zebrafish.

In this work, we report the characterization of the single *Nidogen/entactin* (*Ndg*) gene in *Drosophila.* We found that *Ndg* was highly expressed in the *Drosophila* embryo and NDG protein was abundant in all basement membranes (BMs) suggesting an essential function during development. In contrast, analysis of *Ndg* mutants revealed that loss of NDG did not affect viability and fertility in general, arguing for a negligible role during development. This was further confirmed by examination of BM assembly in *Ndg* mutant embryos revealing no overt changes in overall BM protein distribution. Interestingly, ultrastructural SEM analysis and permeability assays revealed a porous BM surface and severe defects in BM barrier function accelerating the disruption of imaginal disc BMs under osmotic stress conditions. Moreover, *Ndg.* mutant larvae displayed a range of strong behavioral phenotypes such as reduced responses to external stimuli, impaired motility and climbing performance as well as altered crawling behavior and gravitaxis. Our morphological analysis of the larval nervous system indicated that these phenotypes correlated with improper positioning of neurons, defasciculation of nerves as well as severe defects in chordotonal organ and neuromuscular junction (NMJ) organization. We also found that *Ndg* genetically interacted with the *Leukocyte-antigen-related-like* (*Lar*) receptor gene to promote NMJ differentiation and plasticity. Taken together, our analysis of *Drosophila Ndg* did not support an essential role of NDG for the assembly of BMs but suggested that NDG essentially contributes to the robustness of the BM barrier and ensures ECM plasticity in response to external cues.

## Results and Figures

### *Ndg* transcript expression and protein localization

Developmental Northern blot analysis revealed high levels of *Ndg* expression in
*Drosophila* embryos after gastrulation (Sup. Fig. 1). We therefore applied fluorescence *in situ* hybridization on *white*^*1118*^ (*w*^*1118*^) embryos to analyze *Ndg* mRNA distribution in comparison to NDG protein localization revealed by antibody staining (Fig. 1). In line with our Northern blot analysis, *Ndg* expression was apparent at stage 11/12 and could be detected in single cells of the head, especially in the gnathal segments (Fig. 1A, asterisk) as well as in segmentally located patches of cells in the dorsal mesoderm (arrowhead, Fig. 1A). Moreover, the midline-associated, mesodermal dorsal median cells (DMC, Fig. 1B, C) as well as four surrounding somatic myoblasts (sMB, Fig. 1B) exhibited strong *Ndg* expression whereas we detected only very weak signals in the amnioserosa (AS, Fig. 1C). At this time NDG protein, embedding DMCs and sMBs, formed a thin sheet between ecto-and mesoderm and outlined the region of the prospective anal plate. After germ band retraction (Fig. 1D) mRNA expression had increased and was now visible in the forming dorsal and ventral muscles, the amnioserosa (AS) and the segmentally located chordotonal organs (Cho). This was in agreement with the expression profile obtained from developmental Northern blot analysis which revealed a strong *Ndg* mRNA increase in older embryos (Sup. Fig. 1). Internal views indicated a decrease of expression in the DMCs (arrow) whereas *Ndg* was now expressed in the esophageal visceral muscle primordium (eVM, arrowhead) and the joint region between hind-and midgut (Fig. 1E). At this time, accumulation of NDG protein in the forming BMs around the developing brain and ventral nerve cord (VNC), the differentiating tracheal system, the future digestive tract as well as the forming somatic muscles was observed (Figure 1D, E). Embryos at stage 16 displayed strong mRNA expression in somatic and visceral muscles and in the cap cells (CC in Figure 1H) of the chordotonal organs, whereas the protein localized to all embryonic BMs (Figure 1F-I). Interestingly, whereas *Ndg* mRNA expression was absent from neuronal tissues, the BMs surrounding the chordotonal neurons (CN), the VNC and the brain were highly NDG-positive (Figure 1H, I). In summary, *Ndg* mRNA expression during embryogenesis followed a dynamic and distinct pattern involving mostly mesodermal cells and tissues but also chordotonal cap cells and the amnioserosa whereas the protein was highly abundant in all embryonic BMs. Interestingly, no transcript expression was observed in hemocytes or fat body cells although these tissues strongly express other BM components like Laminin or type IV Collagen (Le Parco et al., 1986; Pastor-Pareja and Xu, 2011; Rodriguez et al., 1996; Wolfstetter and Holz, 2012).

**Figure 1:**
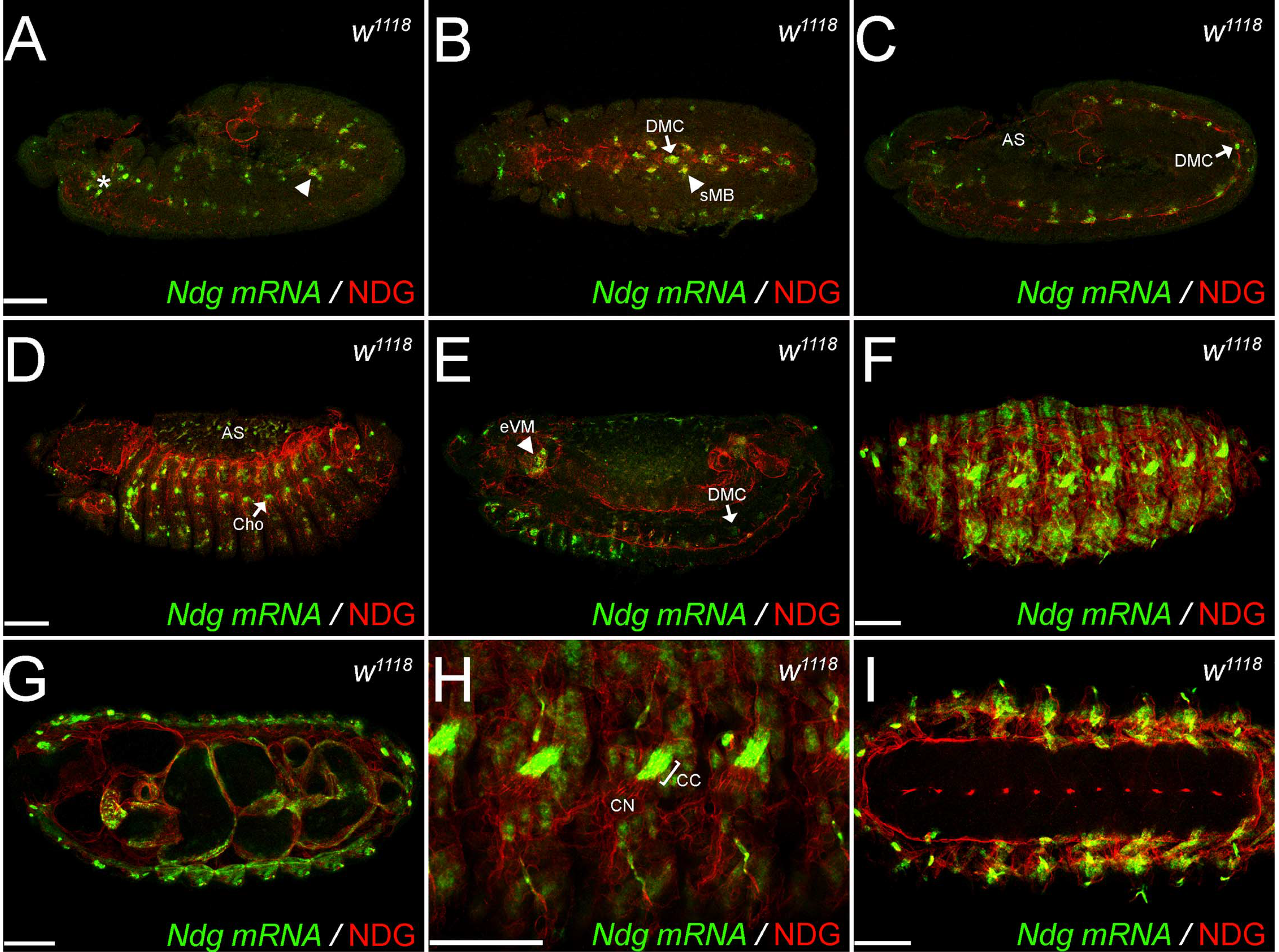
*Ndg* transcript expression and protein localization during embryonic development. Confocal stacks of *Drosophila white* (w^1118^) embryos at developmental stage 12 (**A-C**), stage 14 (**D, E**) and stage 16 (**F-I**). *Ndg* mRNA (green) was visualized by fluorescence *in situ* hybridization (FISH), and antibody staining against NDG (red) reveals the secreted protein. Embryos are orientated in lateral views except for B, G and I which represent ventral views. AS: amnioserosa, CC: chordotonal cap cell, Cho: chordotonal organ, DMC: dorsal median cell, eVM: esophageal visceral mesoderm, CN: chordotonal neuron, sMB: somatic myoblast. Asterisk and arrowhead in (**A**) depict *Ndg* expression in the head region and the dorsal mesoderm respectively. Scale bars = 50 μm.

### Localization of NDG to BMs depends on Laminin

We next asked whether the absence of other major BM components affects the incorporation and localization of NDG within the BM. Therefore we performed NDG antibody staining on late stage 16 embryos lacking either Laminin, type IV Collagen or Perlecan (Figure 2). Control siblings exhibited strong NDG staining of all embryonic BMs and around the chordotonal organs (Figure 2A). To differentiate between the NDG localization properties of both Laminin trimers we employed either *LanA* or *wing blister* (*wb*) loss of function mutations that impair formation and secretion of only one Laminin trimer (Urbano et al., 2009). Interestingly, we found a strongly disrupted and punctuated NDG pattern in transheterozygous *LanA*^*9-32*^/*Df LanA* embryos lacking the Laminin A-yielding trimer (Figure 2B), whereas we did not detect changes in *wb*^*HG10*^/*Df wb* embryos in which the Wb-containing trimer is absent (Figure 2C). The loss of all secreted Laminin trimers in embryos lacking the only Laminin γ-subunit (*LanB2*^knod^/*Df LanB2*) resulted in a dramatic loss of NDG localization (Figure 2D) indicating a redundant function of both Laminin trimers for the localization of NDG to BMs. On the other hand, embryos deficient for *trol* (the Perlecan encoding locus) as well as embryos carrying a deficiency for the two *Drosophila* type VI Collagens *(Col4a1* and *viking)* displayed no changes in NDG localization and distribution (Figure 2E, F) suggesting that Laminin is the primary crucial component for NDG distribution and localization to BMs.

**Figure 2:**
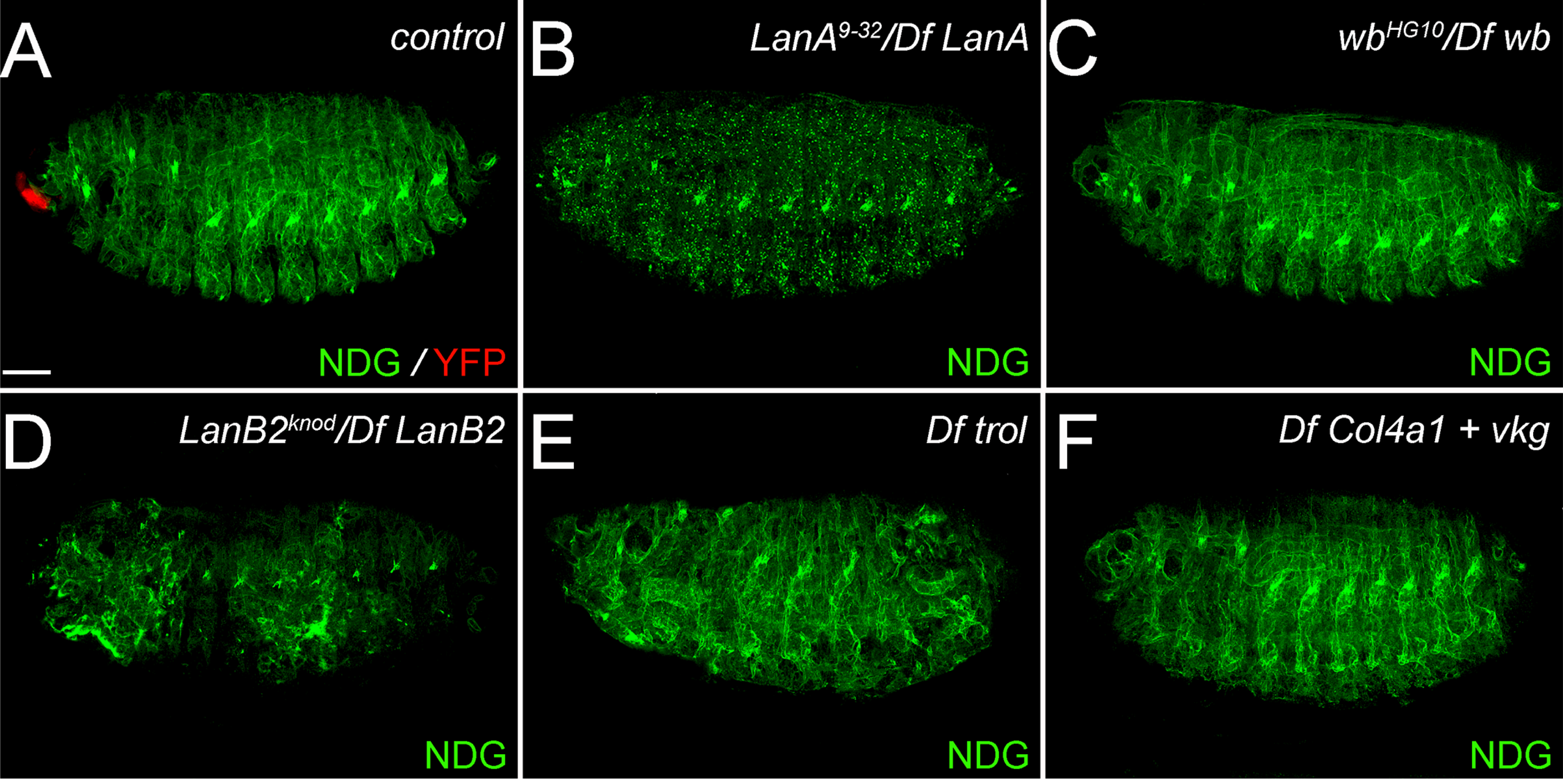
NDG localization in the absence of core BM proteins. Stage 16 embryos in lateral orientation were stained for NDG protein (green). (**A**) Control sibling embryo with balancer-associated YFP expression (**red**) displays proper NDG localization to BMs with strong accumulation at the chordotonal organs. (**B**) Punctate NDG pattern in a *LanA^9-32^JDf LanA* transheterozygous embryo. (**C**) NDG protein distribution appears unaffected in *wb^HG10^JDf wb* embryos. (**D**) Disruption of the NDG pattern in *LanB2^knod^JDf LanB2* embryos. (**E**) *Df trol* (loss of Perlecan) embryos display a segmentation phenotype due to the additional loss of the adjacent *giant* locus but no changes in NDG localization. (**F**) Regular NDG distribution in an embryo deficient for the two *Drosophila* type VI Collagen encoding loci *Col4a1* and *viking (vkg).* Scale bar = 50 μm.

### Generation of *Ndg* deletion mutants

In order to investigate the function of *Ndg* in *Drosophila*, we employed imprecise excision of the Minos transposon insertion *Mi{ET1}MB04184* to generate deletions in the *Ndg* locus (Sup. Fig 2A). Two excision lines were obtained in which 0.4 kb respectively 1.4 kb of the *Ndg* locus including parts of the 5’ UTR were deleted (henceforth referred to as *Ndg*^*Δ0.4*^. and *Ndg*^*Δ1.4*^, see also Material and Methods). Further analysis employing NDG-specific antibodies revealed that the strong NDG staining observed in control siblings (Sup. Fig 2B) was completely absent in homozygous *Ndg*^*Δ1.4*^. embryos (Sup. Fig 2C), whereas faint NDG expression could be detected in the *Ndg*^*Δ0.4*^. mutant (Sup. Fig 2D). Therefore, we concluded that the deletion in *Ndg*^*Δ1.4*^. resulted in a protein null allele while the smaller molecular lesion in *Ndg*^*A0A*^. caused a hypomorphic *Ndg* allele characterized by strongly reduced NDG expression.

### Phenotypic analysis of *Ndg* mutants

*Nidogen* mutants generated in *C. elegans* and mouse are viable and display only mild phenotypes (Bader et al., 2005; Bose et al., 2006; Dong et al., 2002; Kang and Kramer, 2000; Murshed et al., 2000; Schymeinsky et al., 2002), a surprising finding which is in strong contrast to the essential developmental roles demonstrated for the other BM molecules (Arikawa-Hirasawa et al., 1999; Clay and Sherwood, 2015; Poschl et al., 2004; Yao, 2017). In agreement with these former observations, *Drosophila Ndg*^Δ^ mutants were viable and fertile and could be maintained as homozygous stocks. Despite the lack of any obvious developmental function we noticed several peculiarities in *Ndg*^*Δ*^ homozygous stocks that were more apparent in the *Ndg*^*Δ1.4*^. line when compared to *Ndg*^*Δ0.4*^. In non-crowded, standard culture conditions, *Ndg*^*Δ*^ pupal cases were preferentially formed in the lower half of the vial or even directly on the food, indicating potentially impaired larval climbing abilities (Sup. Figure 3A, B). We also noticed a difference in the orientation of the pupal cases which was shifted towards the horizontal axis in *Ndg*^*Δ*^ vials (Sup. Fig 3B). Moreover, we observed a decrease in fecundity in homozygous *Ndg*^*Δ*^ animals compared to the *w*^*1118*^. control (Sup. Fig 3C) and *Ndg*^*Δ1.4*^. flies additionally exhibited incompletely inflated wing blades at low (~5 %) penetrance rates (Sup. Fig 3D, E).

### Distribution of major BM components is not altered in *Ndg* mutant embryos

To analyze the proposed function of NDG as universal ECM cross linker that connects Laminin and Collagen layers of BMs (Timpl and Brown, 1996), we studied the distribution of known BM components in *Ndg*^*Δ1.4*^/*Df Ndg* embryos. Therefore, we applied antibodies raised against Laminin subunits and trimers, type IV Collagens and Perlecan as well as the Collagen-associated molecule SPARC (Fig 3 and Sup. Fig 4). In addition, we employed GFP-trap insertions in the loci of *viking (vkg::GFP)*, one of the two *Drosophila* type IV *Collagen* genes, and *terribly reduced optic lobes (trol::GFP)*, encoding for *Drosophila* Perlecan (Sup. Figure 4).

**Figure 3:**
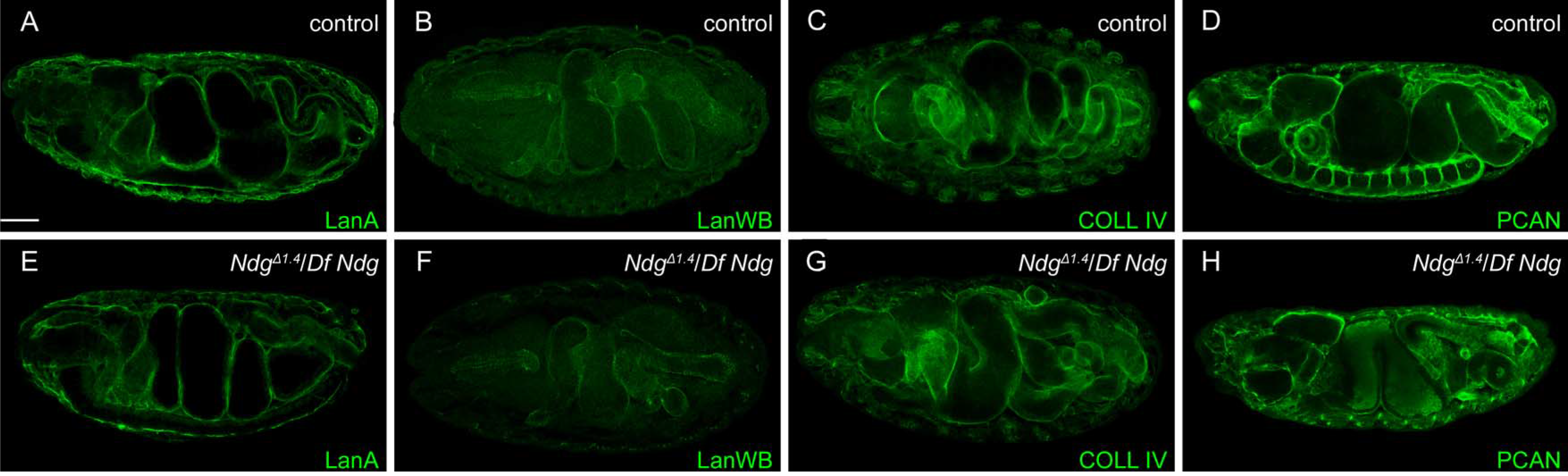
BM formation in the absence of *Ndg.* (**A-D**) Distribution of LanA, LanWb, COLL IV and PCAN in control and transheterozygous *Ndg^*Δ1.4*^IDf Ndg* embryos (**E-H**) at stage 16. Embryos appear in in lateral (**D**), dorsolateral (**A**, **B**, **E**, **F**, **H**) or ventrolateral orientation (**C**, **G**). Scale bar = 50 μm.

In case of Laminin, all antibodies employed stained embryonic BMs indicating reactivity with secreted Laminin (Figure 3A, B). Laminin A (LanA) outlined the BMs of all internal organs in control and *Ndg* mutant embryos (Fig 3A, E), whereas antibody staining against the Laminin α1, 2 subunit Wing blister (LanWb) was found mainly in the BMs around the digestive tract and the apodemes of control and *Ndg* mutant embryos (Fig 3B, E). This was in agreement with antibody staining against secreted K-cell Laminin (LanKc) which appeared similar in *Ndg* mutants compared to control embryos (Sup. Fig 4A, D). Additionally employed antibodies against the Laminin β- and γ-subunits (referred to as LanB1 and LanB2, Sup. Fig 4B, C, E, F) demonstrate BM staining of control and *Ndg* mutant embryos and due to the strong reactivity with non-secreted Laminin intermediates also labeling of hemocytes and fat body cells. Further analyses employing antibodies against Collagen IV (COLL IV, Fig 3C, G), the *vkg::GFP* protein trap (Sup. Fig 4G, J), and the collagen-binding protein SPARC (Sup. Fig 4H, K) revealed that these proteins were present in embryonic BMs of *Ndg* mutants. Finally, we did not observe obvious differences in Perlecan staining of control and *Ndg* mutant embryos detected by either PCAN-specific antibodies (Fig 3D, H) or *trol::GFP* (Sup. Fig 4I, L). Taken together, the complete loss of NDG seems not influence the formation or overall assembly of embryonic BMs.

**Figure 4:**
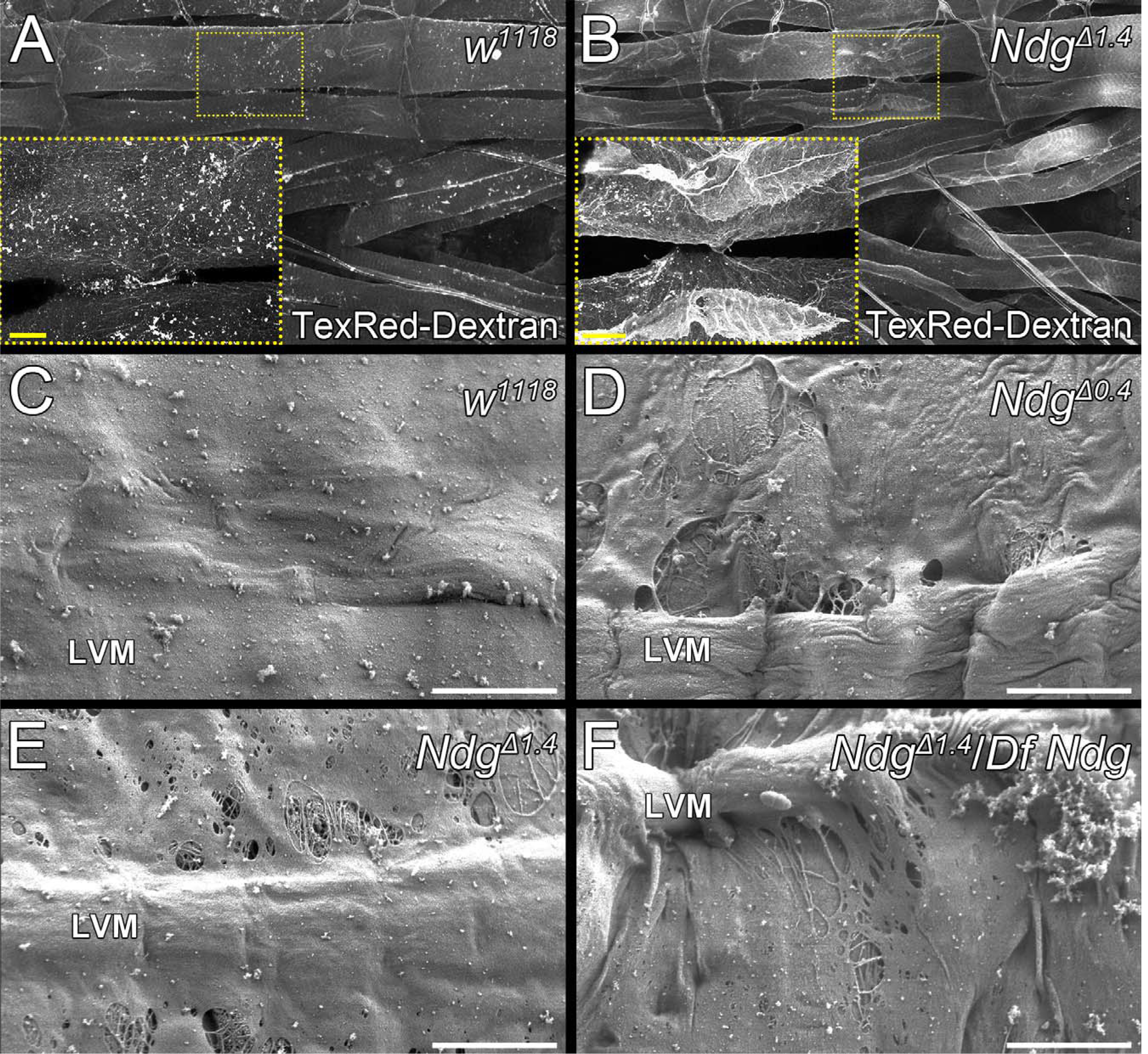
*Ndg* is required for permeability, ultrastructure and mechanical stability of BMs. (**A, B**) Confocal sections from somatic muscle preparations of control (**A**) and *Ndg*^*Δ1.4*^ (**B**) larvae incubated with low weight TexasRed-coupled Dextran (TexRed-Dextran, white) to reveal BM leakage. Insets are higher magnifications of the highlighted areas. Confocal images were acquired with identical channel settings. (**C-F**) Visceral BM ultrastructure of dissected *w^1118^.* control (**C**) 3^rd^ instar larvae and homozygous (**D, E**) and transheterozygous (**F**) *Ndg* mutants revealed by SEM. LVM: Longitudinal visceral muscle. Scale bars = 20 μm (A, B), 5 μm (**C-F**).

### Loss of NDG affects BM ultrastructure and function

The viability of *Ndg*^*Δ*^ mutant flies and the presence of all BM proteins analyzed so far (Fig 3 and Sup. Figure 4) suggested that BM assembly was not severely affected by the absence of NDG, therefore questioning its proposed role as universal ECM linker molecule. However, the loss of NDG might impair BM stability and function. To address this in more detail we dissected *w*^*1118*^. and *Ndg*^*Δ1.4*^. 3^rd^ instar larvae and employed a Dextran-based permeability assay in addition to SEM ultrastructural analysis (Figure 4). Spot-like Texas Red (TexRed)-coupled Dextran plaques were observed across the surface of the somatic muscles of *w*^*1118*^ control larval filets (Figure 4A). In comparison, *Ndg*^*Δ1.4*^. larval filets appeared much brighter after dextran staining and we observed a strong leakage of TexRed-Dextran into the muscle proper (Figure 4B) indicating enhanced diffusion through the BM tissue barrier in the absence of NDG.

Further ultrastructural analysis of *w*^1118^ control 3*rd* instar larvae revealed that the visceral BM which covers the surface of the larval midgut appeared as smooth sheet spread across the underlying longitudinal and circular visceral muscles (Figure 4C). In comparison, the surface of the visceral BM in homozygous *Ndg*^*Δ*^ mutant larvae exhibited holes (Figure 4D-F), which were frequently found in close vicinity to the longitudinal visceral muscles (LVM) but also appeared in other areas. We also observed a variation in phenotypic strength between individual larvae. While most null mutant *Ndg*^*Δ1.4*^. animals displayed strong BM phenotypes, a minor fraction also exhibited weaker defects resembling *Ndg*^*Δ0.4*^. larvae. On the other hand, BM defects observed in some of the analyzed *Ndg*^*Δ0.4*^. larvae appeared similar to the null allele. Interestingly, examination of heterozygous *Ndg*^Δ^/*CyO* larvae (Sup. Figure 5B-D) already revealed a weak BM phenotype characterized by few and small surface holes which was never observed in *w*^*1118*^. (Figure 4C) or in additional control larvae from the *w*;* Kr^*If-1*^/*CyO* balancer stock (Sup. Fig 5A), indicating a dose-dependent effect of NDG on BM morphology.

### Loss of *Ndg* leads to BM disruption under osmotic stress

To further investigate whether the absence of NDG comprises the mechanical stability of the BM, we exposed wing imaginal discs to an osmotic shock by incubating them in deionized water. When water diffused into the hypertonic interior of the imaginal discs it created balloon-like swelling of the disc before eventual bursting. Imaginal discs of larvae resembling the genetic background of balanced and homozygous *Ndg*^*Δ*^. animals (*w*^*1118*^ and *w*; Kr*^*If-1*^/*CyO*) withstand the resulting osmotic pressure for ~430 s respectively ~388 s before bursting. In case of wing discs derived from balanced, heterozygous *Ndg*^*Δ*^ animals we observed a slight but not significant increase *(Ndg^*Δ0.4*^/CyO* = 472 s, *Ndg*^*Δ0.4*^.*/CyO* = 520 s; and *Df Ndg/CyO*= 407 s). In contrast to this *Ndg*^*Δ0.4*^. wing discs burst after ~213 s and *Ndg*^*Δ1.4*^. or *Ndg^*Δ1.4*^/Df Ndg* discs tore after ~240 s indicating decreased BM stability (Figure 5).

**Figure 5:**
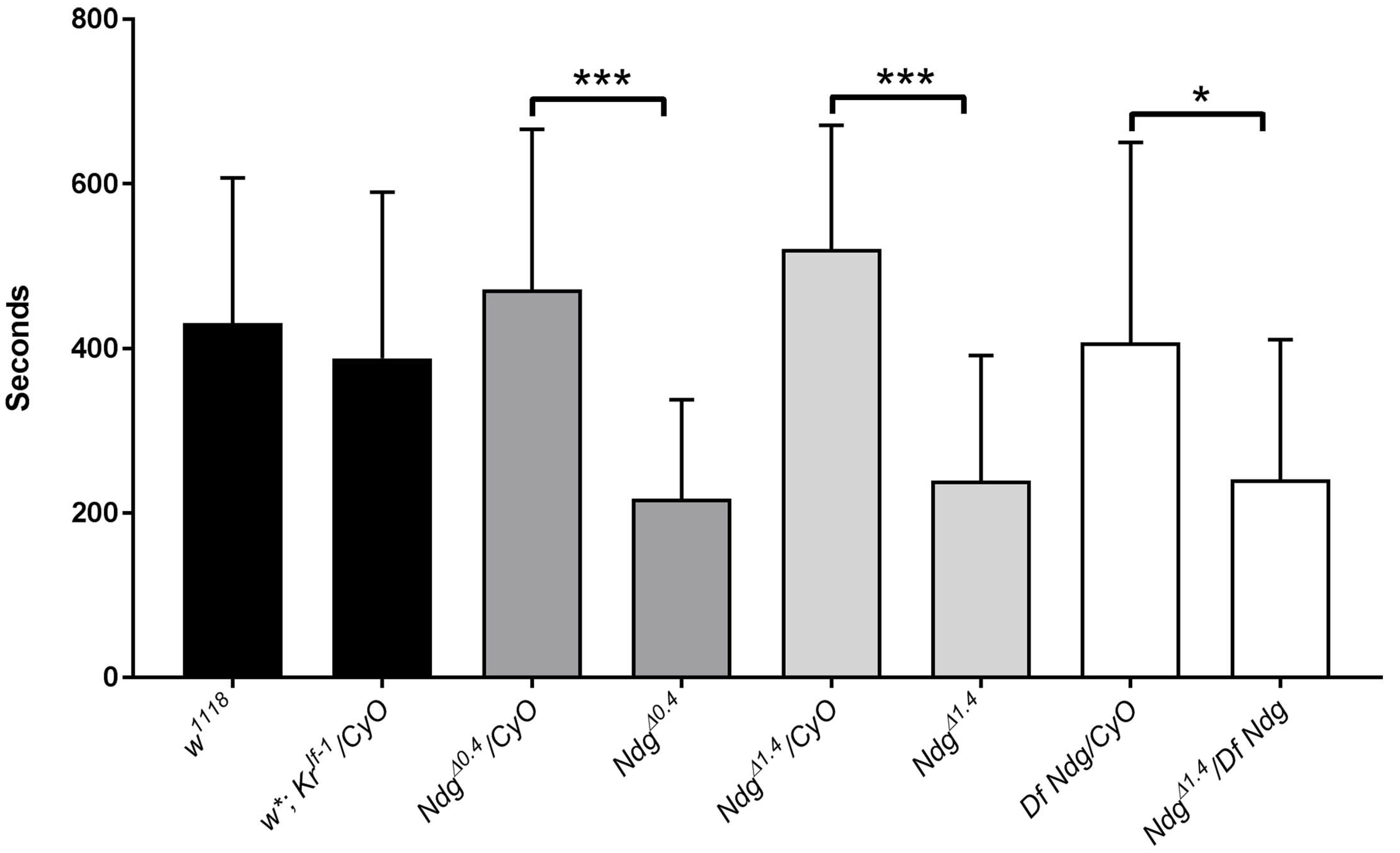
Loss of NDG affects BM stability under osmotic stress conditions. Quantification of osmotic stress applied to 3^rd^. instar larval wing discs of the indicated genotypes. Means and SDs (**whiskers**) are shown. Compared to *w^1118^.* and *w*; Kr*^If-1^/*CyO* balancer controls, imaginal discs from *Ndg* mutant animals resist osmotic stress for a significantly reduced mean time (*Ndg*^*Δ0.4*^/*CyO* = 472 s vs. *Ndg*^*Δ0.4*^. = 213 s, *Ndg^*Δ1.4*^ICyO* = 520 s vs. *Ndg*^*Δ1.4*^., 239 s; and *Df Ndg/CyO* = 407 s vs. *Ndg^A1A^/Df Ndg*, 241 s, n ≥ 20 wing imaginal discs). 1-W anova indicated significant differences among means (F(._7,188_) = 11.8, p < 0.001) and unpaired, two-tailed t-tests were applied to compare heterozygous controls and homozygous samples (***p < 0.001,*p = 0.017).

### Altered crawling behavior and NMJ morphology in *Ndg* mutant larvae

When placed on agar plates supplemented with yeast paste as food source *Ndg*^*Δ1.4*^ 1^st^ instar. larvae were often observed outside the food and exhibited overall lethargic and reduced crawling movements (Figure 6A, B). To monitor and compare their behavioral repertoire we recorded age-matched *w*^*1118*^ controls and *Ndg*^*Δ1.4*^ larvae that were placed on Agar plates in the absence of food (Fig 6C, D). Crawling of control *w*^*1118*^ larvae was dominated by smooth forward movement with occasional turns and pauses that allowed the larvae to explore the whole experimental arena (Figure 6C, E and Sup. Movie). In contrast, *Ndg* mutants generally moved within a much smaller area and displayed a variety of crawling defects including excessive head turning as well as exaggerated rolling and bending motions (Figure 6D, F and Sup. Movie). Interestingly, the phenotypic strength exhibited some variation since 70 % of 2^nd^ instar larvae (n > 40) and half of the analyzed 3^rd^ instar larvae (n > 40) displayed rather strong behavioral abnormalities whereas the remaining animals exhibited weaker crawling defects. We further analyzed crawling of *Ndg*^*Δ*^ 3^rd^ instar larvae in more detail, investigating the velocity and the stride frequency of the undisturbed crawling pattern (Figure 6G, H). Comparison of the mean crawling velocity and the stride frequency showed comparable differences between the analyzed genotypes. The mean crawling velocity of *Ndg*^*Δ0.4*^ mutant larvae was slightly but not significantly reduced whereas *Ndg*^*Δ1.4*^ larvae showed a significantly reduced mean velocity compared to *w*^*1118*^ larvae (Figure 6G). Also the stride frequency in both *Ndg*^*Δ*^ strains was significantly lower when compared to w larvae (Figure 6H). However, *Ndg*^*Δ*^ mutants displayed the more severe phenotype in terms of reduced crawling speed and stride frequency (Figure 6G, H).

**Figure 6:**
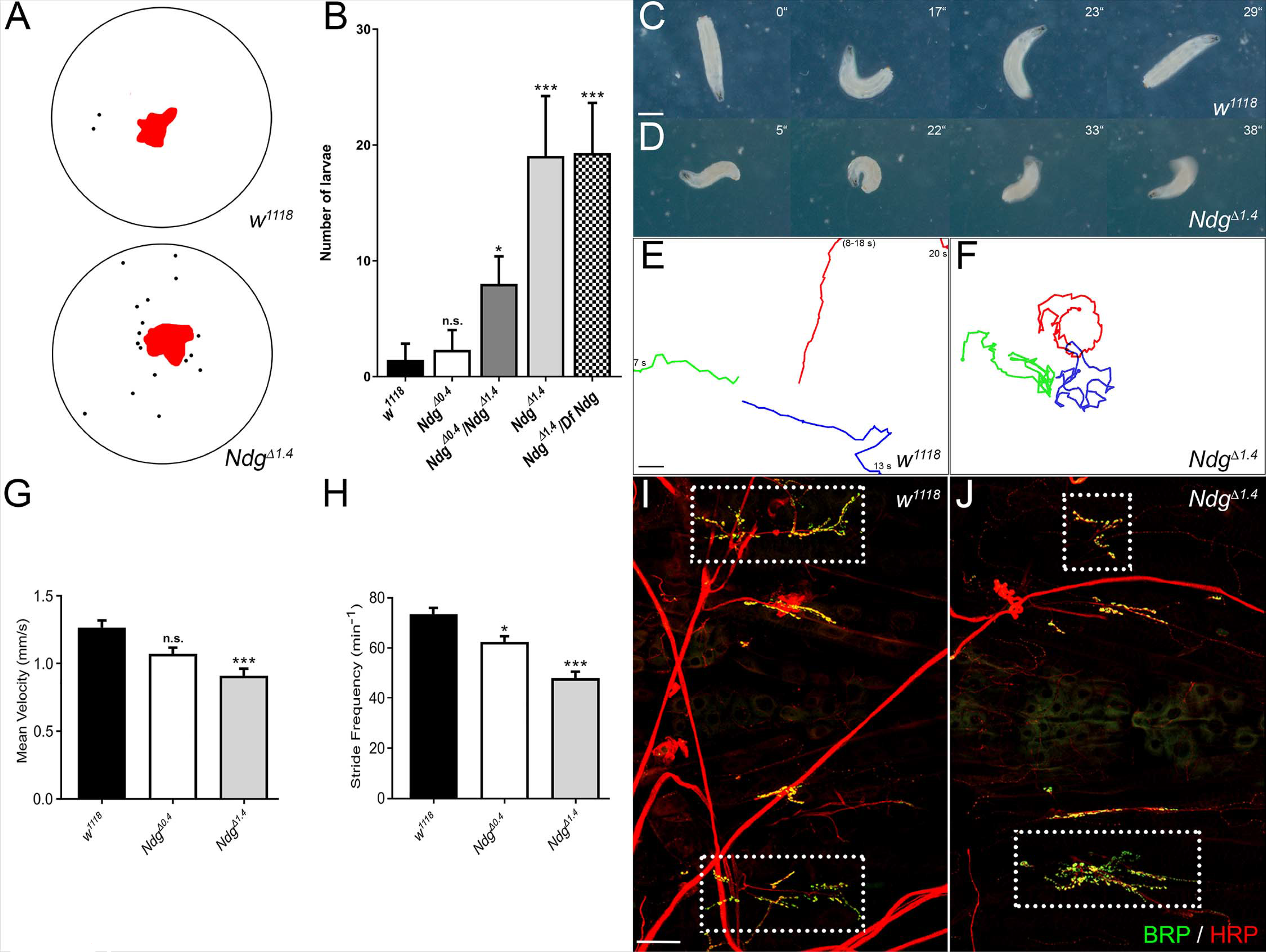
Altered crawling behavior and neuromuscular junction phenotypes of *Ndg* mutant larvae. (**A**) Schematic example of the observed distribution of *w*^1118^.and *Ndg*^*Δ1.4*^. 1^st^ instar larvae (black dots) on agar plates (depicted as circles) outside a central food source (**red**). (**B**) Quantification of the larval distribution assay. Mean of larvae outside the food: *w*^*1118*^ = 1.44, *Ndg^*Δ0.4*^* = 2.31, *Ndg^*Δ1.4*^* = 19.1, *Ndg^*Δ0.4*^/Ndg^*Δ1.4*^ = 8, Ndg^*Δ1.4*^/Df Ndg* = 19.3; 1-W anova p < 0.001, Tuckey’s multiple comparisons test with adj. p-value: *w*^1118^ versus *Ndg*^*Δ0.4*^ = 0.947 non-significant (n.s.), *w* versus *Ndg^*Δ0.4*^/Ndg^*Δ1.4*^*. = *p < 0.01, *w^1118^.* vs. *Ndg*^*Δ1.4*^. ***p < 0.001, *w^1118^.* vs. *Ndg^A14^/Df Ndg =* ***p < 0.001, n = 400 larvae for each genotype. (**C, D**) Snapshots taken from crawling recordings of *w*^*1118*^. and *Ndg*^*Δ1.4*^. 2^nd^ instar larvae. (**E, F**) Representative tracking patterns of crawling *w*^*1118*^. and *Ndg*^*Δ1.4*^. 2^nd^ instar larvae. Individually colored lines represent recordings of 25 s. Time points are denoted if the larva left the monitored area before the end of recording. Time intervals in brackets indicate absence from the recorded area. (**G**) Quantification of mean crawling velocity measured as 5s of uniform crawling from recordings of *w*^*1118*^. and *Ndg*^*Δ1.4*^. L3 larvae. Mean: *w*^*1118*^ = 1. 265, *Ndg*^*Δ0.4*^ = 1.07, *Ndg^*Δ1.4*^* = 0.907 mmJs) (K-W anova p < 0.0001, Dunn’s-test with mult. adj. p-value: *Ndg^*Δ0.4*^* = 0.0597, *Ndg*^*Δ1.4*^. ***p < 0.0001 versus w *^1118^.* (**H**) Quantification of the stride frequency calculated from 5 strides in a row from recordings of *w*^*1118*^. and *Ndg^*Δ1.4*^* 3^rd^. instar larvae. Mean: *w*^*1118*^ = 73.32, *Ndg^*Δ0.4*^* = 62.25, *Ndg^*Δ1.4*^* = 47.75 strides/min.; K-W anova p < 0.0001, Dunn’s-test with mult. adj. p-value: *Ndg^*Δ0.4*^*.*p = 0.0207, *Ndg*^*Δ1.4*^. ***p < 0.0001 versus *w*^*1118*^. Error bars represent SEM. N >23 larvae for each genotype (**I, J**) Neuromuscular junctions (NMJs, outlined by dashed boxes) innervating muscles 6J7 in the 2 ^nd^ larval abdominal segment of *w^1118^.* and *Ndg^*Δ1.4*^* 3^rd^. instar larvae are stained with anti-Bruchpilot (BRP, green) and anti-HRP (**red**). Scale bars = 1 mm (C, E), 50 μm (**I**).

These crawling phenotypes prompted us to examine neuromuscular junction (NMJ) morphology in wandering 3^rd^ instar larvae. We therefore dissected larval filets and stained them with anti-Bruchpilot (BRP) antibodies to label T-zones in the synaptic boutons and employed anti-HRP staining to reveal the overall NMJ morphology (Figure 6I, J). We focused on the NMJ at muscle 6/7 in the 2^nd^ abdominal segment because it covers a considerably large area, innervates two muscles, and contains many synaptic boutons. While these NMJs in *w*^1118^ control larvae appeared to be equally sized, we observed shape and size differences of the corresponding NMJs in *Ndg*^*Δ*^ mutants, even within the same animal (Fig 6J). We also noticed a higher degree of synaptic branching and increased bouton density (Figure 6J and 8B, C) which appeared to be independent of the NMJ size. Taken together, these results suggest that loss of NDG leads to defects in NMJ organization presumably resulting in the climbing and crawling defects observed in *Ndg*^*Δ*^ mutant larvae.

### Loss of NDG leads to reduced reaction to vibrational stimuli and aberrant chordotonal organ morphology

Our previous analyses revealed that developing chordotonal organs highly expressed *Ndg* in their cap cells and eventually became embedded in a NDG-positive BM. Notably, some of the larval crawling defects and the altered orientation of pupal cases observed in the absence of NDG could be interpreted as a result of impaired proprioception. Therefore, we wanted to characterize chordotonal function in *Ndg*^*Δ*^ mutants on a behavioral level and additionally analyze larval PNS morphology with respect to the chordotonal organs (Figure 7). Stimulus-induced, relative body length reduction upon applied vibrational stimuli was compared between 3^rd^ instar *w*^*1118*^ control and *Ndg*^*Δ*^ larvae (Figure 7A, B). The body length reduction of control larvae in response to vibrational stimuli was on average ~14% whereas homozygous *Ndg*^*Δ0.4*^ and *Ndg*^*Δ1.4*^. larvae showed significantly reduced retraction values (Fig 7B). *Ndg^A1A^* mutants again displayed the most severe phenotype with a nearly abolished reaction (~4% body length reduction) to vibrational stimuli (Figure 7B).

**Figure 7:**
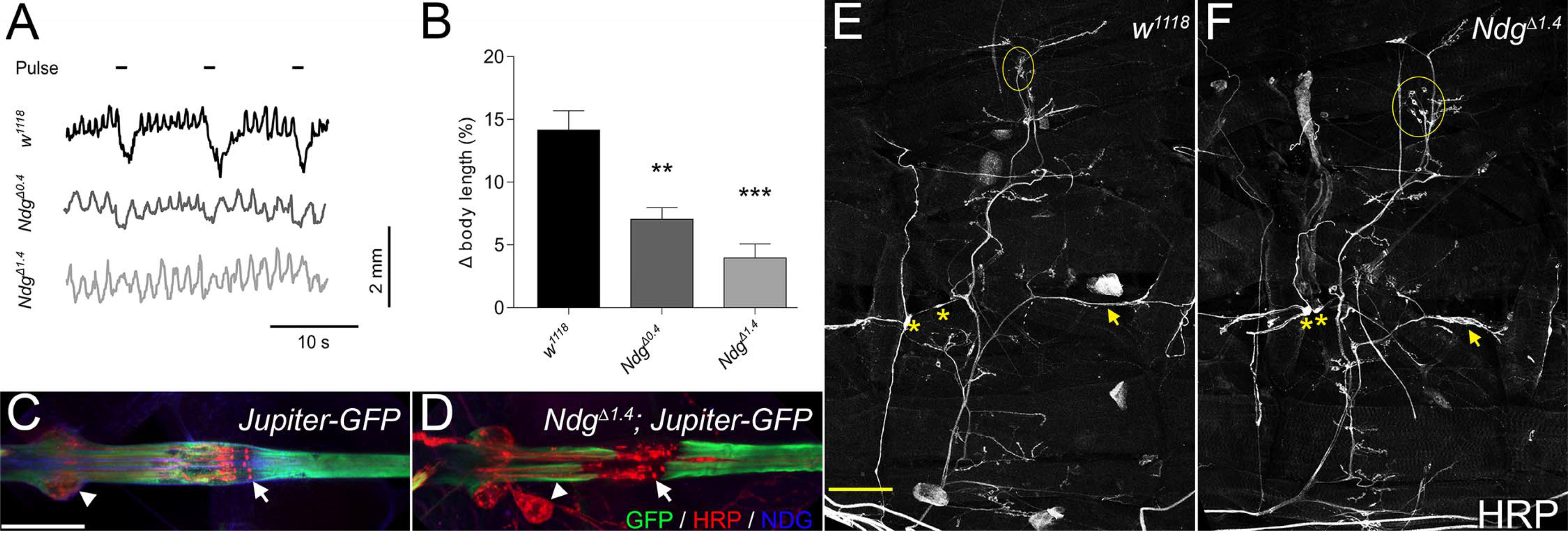
Loss of *Ndg* affects response to vibrational stimuli and PNS morphology. (**A**) Body length recordings of *w*^*1117*^control and *Ndg*^*Δ0.4*^ and *Ndg* mutant larvae exposed to pulsed vibrational stimuli (indicated by black bars). (**B**) Quantification of body length reduction induced by vibrational stimulation in *w*^*1118*^ control and *Ndg* mutant larvae. The relative body length reduction (A body length) is calculated from the mean larval length before and during vibrational stimulation. Error bars represent SEM, n = 24, **p < 0.01 ***p < 0.001 versus w^1118^. K-W anova p < 0.0001 and Dunn’s-test with mult. adj. p-value was applied for statistical analysis. *Ndg^*Δ0.4*^* vs. w^1118^.p = 0.0061, *Ndg*^*Δ1.4*^. p < 0.001 vs. *w^1118^.*.(**C, D**) Morphology of the lateral pentascolopidial chordotonal organ (lch5) of control and *Ndg^*Δ1.4*^* larvae revealed by Jupiter::GFP fusion protein (**green**) expression, NDG-(**blue**), and HRP-(**red**) antibody staining. Arrows highlight the sensory cilia whereas neuronal cell bodies are marked by arrowheads. (**E, F**) HRP labeling was employed to reveal the nervous system in one hemisegment of control and *Ndg^A1′^* ^4^ mutant larvae. Structures of the peripheral nervous system (**PNS**) such as ddA neurons (surrounded by circles) transmitting nerves (**arrow**) and neuronal cell bodies (**asterisks**) are highlighted. Scale bars = 50 μm (**C**), 100 μm (**E**).

To reveal potential morphological changes underlying the observed behavioral phenotypes, we examined the chordotonal organs of 3^rd^ instar larvae. Therefore, we utilized the *Jupiter::GFP* fusion protein which localizes to the microtubule network (Karpova et al., 2006). In addition, we employed anti-NDG and horseradish peroxidase (HRP) staining (Jan and Jan, 1982) to reveal the chordotonal BM and associated neurons respectively (Fig 7C, D). A NDG-positive BM surrounded the whole mechanosensory organ of *Jupiter::GFP* control larvae including the bulging cell bodies of the chordotonal neurons (Fig 7C, arrowhead) which closely aligned to the scolopale cell. A NDG-positive sheet also surrounded the row of aligned sensory cilia (Fig 7C, arrow). In contrast to this regular alignment, *Ndg*^*Δ1.4*^. mutant larvae exhibited detachment of the chordotonal neurons and dislocation of their cell bodies (Fig 7D, arrowhead) from the scolopale cell as well as disrupted alignment of the sensory cilia (Fig 7D arrow). We noticed that the penetrance of these phenotypes varied even between the chordotonal organs of corresponding hemisegments in a single larva (Sup. Figure 6B, C). However, we did not observe chordotonal organs in *Ndg* mutant larvae that resembled wild-type morphology (Sup. Fig 6A).

In addition to the chordotonal organ defects we noticed aberrations in other parts of the larval peripheral nervous system (Fig 7F). The cell bodies of the dorsal dendritic arborization sensory neurons (ddA neurons) failed to cluster in *Ndg^*Δ1.4*^* mutant larvae (Fig 7F circles). Moreover, transmitting nerves frequently exhibited defasciculation (Fig 7E and F arrow) and positioning of neuronal cell bodies (Fig 7F asterisks) was often altered when compared to the controls (Fig 7E asterisks).

### *Ndg* interacts genetically with the *Lar* receptor at the larval NMJ

The differences in shape and size between corresponding NMJs 6/7 in the 2^nd^ abdominal segment and the higher degree of synaptic branching in *Ndg*^*Δ1.4*^ mutants (Figure 7E, F) prompted us to analyze this phenotype in more detail with regard to potential genetic interaction partners. Upon closer examination of the NMJ geometry in *Ndg*^*Δ*^ mutant larvae we observed that in contrast to wild-type NMJs (Figure 8A) the overall branching was increased and major branching points were shifted towards the center of the junction (Fig 8B). We also observed bouton clustering and a significantly enhanced number of boutons per square at *Ndg*^*Δ*^ junctions suggesting that NDG actually suppresses NMJ maturation (Fig 8C). Notably, these defects were found to be independent from the variable NMJ size observed in *Ndg*^*Δ*^ animals. Moreover, heterozygous *Ndg*^*Δ1.4*^/*CyO* NMJs exhibited increased bouton clustering and a tendency to branch prematurely (Figure 8D) suggesting that these phenotypic characteristics could be modified upon genetic interaction.

Homozygous as well as heterozygous null mutants of the *receptor protein tyrosine phosphatase (RPTP) Leukocyte-antigen-related-like (Lar)* have been described to exhibit a strongly reduced bouton number at the larval NMJ (Kaufmann et al., 2002). In agreement to this we found drastically reduced NMJs in the rarely occurring homozygous *Lar.^13.2^* 3^rd^ instar larvae (data not shown), whereas heterozygous *Lar*^*13.2*^/CyO animals (Fig 8E) exhibited reduced branching and bouton numbers. In double-heterozygous *Lar*^13.2^, *Ndg*^*Δ1.4*^./*CyO* larvae, we observed mutual repression of the phenotypes caused by one copy of either *Lar*.^13.2^ or *Ndg*^Δ^. leading to NMJs that resembled a more “wild-type” morphology (Fig 8F). These results indicate that *Ndg* and *Lar* function together to promote differentiation of the larval NMJ in *Drosophila*.

**Figure 8:**
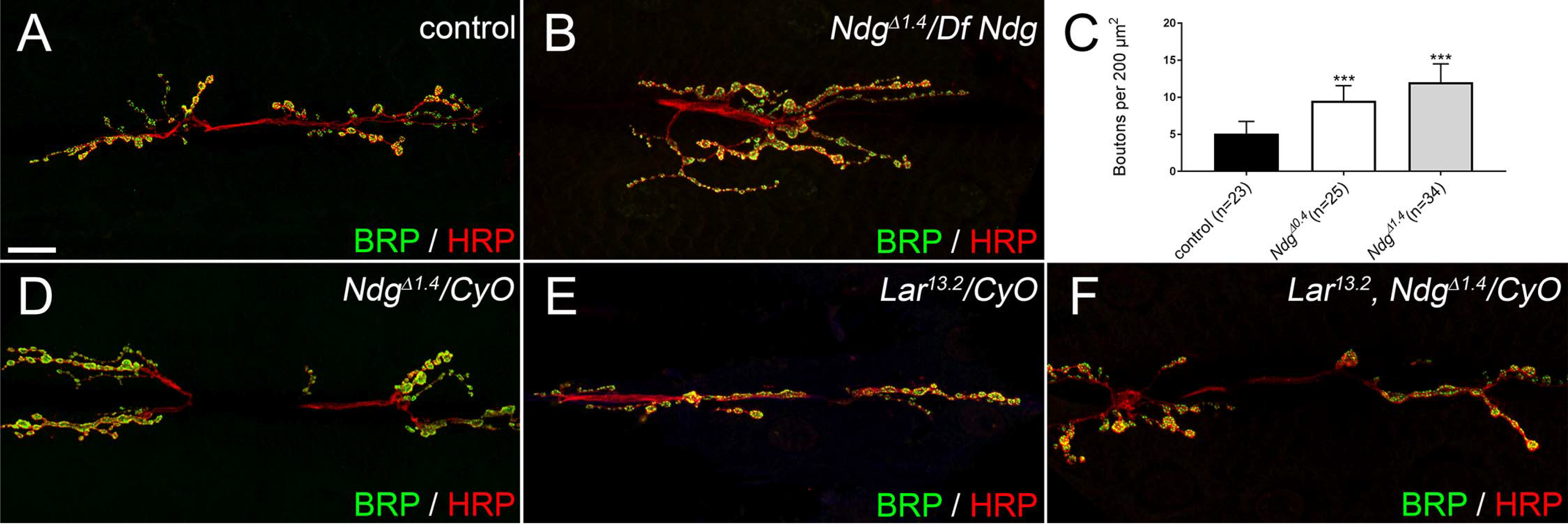
Genetic interaction between *Ndg* and Lar. NMJs of 3^rd^ instar larva stained with anti-Bruchpilot (BRP, green) and anti-HRP (**red**). (**A**) Control NMJ of muscle 6/7 in the 2^nd^ abdominal segment. (**B**) *Ndg.* mutant (*Ndg*^*Δ1.4*^./*Df Ndg*) larvae exhibit increased and premature synaptic branching as well as increased bouton number and clustering. (**C**) Quantification of bouton densities (bouton number per area) of control and *Ndg*^*Δ*^ larvae. (**D**) Heterozygous *Ndg*^*Δ1.4*^/*CyO* larvae display mild branching defects and increased bouton clustering. (**E**) Heterozygous *Lar^13.2^ICyO* NMJ with reduced branching and boutons. (**F**) *Lar*^13.2^, Ndg^A14^/*CyO* larvae exhibit an intermediate phenotype when compared to the single mutations (**D and E**). Scale bar = 20 μm.

## Discussion

### Analysis of *Ndg* expression and NDG protein distribution

In this work we have characterized the single *Nidogen/entactin* gene in *Drosophila melanogaster.* In the developing embryo, *Ndg* expression followed a highly dynamic pattern involving diverse mesodermal cell types like somatic muscle founder cells, mature visceral and somatic muscles as well as dorsal median cells. *Ndg* expression in a subset of somatic muscle founder cells had been previously reported by (Artero et al., 2003) and detailed analysis of an intronic *“Ndg* muscle enhancer” revealed a complex interplay of Forkhead and Homeodomain transcription factors and their binding sites that are required to drive *Ndg* expression in the different mesodermal cell types (Busser et al., 2013; Busser et al., 2012; Philippakis et al., 2006; Zhu et al., 2012). Our analysis further revealed *Ndg* expression in the extraembryonic amnioserosa and in chordotonal cap cells indicating the presence of additional regulatory elements. We did not detect *Ndg* expression in the embryonic CNS but in the closely associated dorsal median cells (DMCs), which express other BM components and ECM receptors such as Perlecan, Dystroglycan, Glutactin as well as Laminin and type IV Collagen (Friedrich et al., 2000; Mirre et al., 1992; Montell and Goodman, 1989; Olson et al., 1990). DMCs provide cues for proper axonal pathfinding during transverse nerve outgrowth and regulate bifurcation of the median nerve in *Drosophila* (Chiang et al., 1994). Therefore, studying a potential role of the secreted ECM in this context would be an interesting topic for future investigations. Interestingly, *Ndg* expression was absent from mesoderm-derived hemocytes and fat body cells although these cells strongly secrete Laminin, type IV Collagens and other ECM components (Le Parco et al., 1986; Pastor-Pareja and Xu, 2011; Rodriguez et al., 1996; Urbano et al., 2009; Wolfstetter and Holz, 2012) indicating that *Ndg* expression is regulated independently from other BM genes.

CEL-Seq transcriptome analysis in *C.elegans* reveals high *nid-1* mRNA expression in mesodermal tissues upon gastrulation (Hashimshony et al., 2015) and NDG proteins strongly associate with body wall muscles in *Drosophila* (this study), *C. elegans*, Ascidians and mice (Ackley et al., 2003; Fox et al., 2008; Kang and Kramer, 2000; Nakae et al., 1993) suggesting an important, evolutionary conserved function in this particular tissue. Indeed, *C. elegans* Nid-1 and NID2 in mice are enriched at neuromuscular junctions and, in agreement with our analyses in *Drosophila*, required for proper NMJ structure and function (Ackley et al., 2003; Fox et al., 2008).

### NDG is not required for BM assembly

Our analysis of NDG distribution in the absence of major BM components revealed a dependency of NDG localization from Laminin but not from type IV Collagen or Perlecan. This is in agreement with studies on γ1III4-mice in which the NDG-binding site in the Laminin γ1 chain had been deleted. This analysis revealed a severe reduction of NDG from BMs as well as perinatal lethality and organ defects similar to those observed in *NIDI*^−/−^. *NID2*^−/−^. double mutant mice which further strengthens the assumption that Laminin plays an essential role in localizing NDG to forming BMs (Bader et al., 2005; Halfter et al., 2002; Willem et al., 2002). The two Laminin heterotrimers in the fly function in a redundant manner to localize NDG to embryonic BMs. In a single loss of function background, however, only absence of the LanA trimer (containing the *Drosophila* Laminin α3/5 homolog) affected NDG localization, suggesting different binding and polymerization activities of the *Drosophila* Laminins. In line with these findings, combined reduction of LanA and NDG around the developing gonad was also observed in βPS-integrin mutant embryos (Tanentzapf et al., 2007). No obvious changes in the distribution of BM core components (Laminins, Collagens and Perlecan) were found in our immunohistochemical analysis after complete loss of NDG, reflecting its non-essential role during development. This finding, further supported by similar analyses in other species (Ackley et al., 2003; Bader et al., 2005; Kang and Kramer, 2000; Kim and Wadsworth, 2000; Rossi et al., 2015) contradicts the proposed role for NDG as linker between inner and outer ECM networks during BM formation, suggesting a negligible role for NDG in overall BM assembly (Ho et al., 2008; Hohenester and Yurchenco, 2013; Jayadev and Sherwood, 2017; Kramer, 2005; Timpl and Brown, 1996; Yurchenco, 2011).

### Loss of NDG affects BM stability and function

Although BM changes are barely detectable in *Ndg*^*Δ*^. mutants by conventional immunohistochemistry, ultrastructural analysis revealed that BM continuity is compromised if NDG is absent or reduced. Our findings that BM barrier function and stability were decreased in *Ndg*^*Δ*^ larvae suggests that NDG, or NDG-mediated crosslinking of other ECM molecules, plays an important role in sealing the assembled BM. Intriguingly, Matsubayashi et al. (Matsubayashi et al., 2017) observe holes in the forming BM around the embryonic *Drosophila* CNS which are progressively closed during hemocyte migration and concurrent Col IV deposition. In addition, studies on mice demonstrate that although Laminin is sufficient for the assembly of early BM scaffolds, incorporation of type IV Collagen is essentially needed for BM integrity and stability (Poschl et al., 2004). Therefore, it is tempting to speculate that crosslinking of hemocyte-deposited Col IV might be impaired in *Ndg* mutants leading to decreased BM stability and barrier function.

The increased number of BM holes in close proximity to the visceral longitudinal musculature in *Ndg* mutants and the reduced tolerance to osmotic shock further implies that NDG protects BM integrity from mechanical stress therefore separating and maintaining different compartments in the larval body. While our experiments demonstrated reduced BM stability in *Ndg*^*Δ*^. larvae, the BM defects observed in the background of different *Ndg* alleles exhibited some phenotypic variation. An explanation for this could be that, although the genetic background of the larvae determines overall BM stability, individual behavior, physical activity, and motility levels could modulate *Ndg* loss of function phenotypes. A comparable finding was made by Tsai and colleagues (Tsai et al., 2012) who reported a process in which larval NMJ size changes in response to LanA levels at the NMJ which are influenced by crawling activity as well as nervous responses to environmental cues. Given the NDG-localizing properties of LanA as well as the variations in NMJ size observed in *Ndg*^*Δ*^ animals, it is intriguing to think of a feedback mechanism that employs spatiotemporal reorganization of the larval BM by NDG to adapt NMJ morphology to altered environmental conditions.

### Loss of NDG affects behavioral responses and PNS morphology

With the exception of *NIDI*^−/−^ *NID2*^−/−^ double mutant mice that die at birth, loss of NDG is generally associated with a range of rather subtle morphological and behavioral phenotypes. Notably, abnormalities described in the absence of NDG are not always fully penetrant or exhibit phenotypic variation (Bader et al., 2005; Bose et al., 2006; Dong et al., 2002; Hobert and Bulow, 2003; Kang and Kramer, 2000; Kim and Wadsworth, 2000; Murshed et al., 2000; Schymeinsky et al., 2002; Zhu et al., 2017). Moreover, subtle behavioral phenotypes can be enhanced by adjusting the experimental conditions (Ackley et al., 2003). In agreement with these analyses, characterization of *Drosophila Ndg*^Δ^. mutants did not reveal an essential developmental function but a range of less overt, often variable phenotypes. After vibrational disturbance of a normal crawling phase, wild-type *Drosophila* larvae show a complex sequence of behavioral pattern. The larvae discontinue their forward movement and show a head retraction (“hunch”) followed by a head turning phase (“kink”) (Bharadwaj et al., 2013; Ohyama et al., 2013; Wu et al., 2011). This sequence of behavior is initiated upon mechanical stimuli delivered to the substrate which are detected by segmental chordotonal organs, mediated by stimulus processing in the central nervous system, and executed under neuromuscular control. Our analysis of the vibrational response behavior in *Ndg*^Δ^. larvae shows a statistically significant reduction in body contraction. This finding points to a crucial role of NDG in vibrational response behavior. However, the precise level of NDG action cannot be inferred from these behavioral data. NDG expression in chordotonal cap cells, the morphological defects observed in larval chordotonal organs as well as the altered geotaxis of *Ndg*^*Δ*^ larvae suggest a contribution to mechanosensation. Defective wing inflation and the aberrant NMJ morphology observed in *Ndg*^*Δ*^ animals additionally indicate that loss of NDG affects the neuromuscular system. Indeed, previous research on *C. elegans* and mouse has uncovered a role of NDG for the structural development of neuromuscular junctions as well as altered locomotor behavior upon loss of NDG (Ackley et al., 2003; Bader et al., 2005; Dong et al., 2002; Fox et al., 2008). Given the high conservation of NDG throughout the animal kingdom (reviewed in (Ho et al., 2008)), this finding could hint at a possible comparable function for NDG in insects.

The receptor protein tyrosine phosphatase (RPTP) Leukocyte-antigen-related-like (Lar) and its cytoplasmic binding partner Liprin-α are required for proper synaptic morphology at the larval NMJ in *Drosophila* (Kaufmann et al., 2002). Interestingly, a specific splice form of LAR has been shown to bind a Laminin/Nidogen-containing protein complex in Hela cells and LAR-like RPTPs genetically interact with *nid-1* to promote presynaptic differentiation in *C. elegans* (Ackley et al., 2005). Our analysis also reveals that a genetic interaction between *Lar* and *Ndg* is conserved at the larval NMJ in *Drosophila.* However, in contrast to the observations in *C. elegans* both factors affected overall NMJ morphology and acted antagonistically in this process.

In conclusion, our initial characterization of *Ndg* mutants in *Drosophila* neither revealed an essential developmental function nor did it support the proposed role for NDG as universal ECM-linker molecule. However, NDG appears to be important for BM sealing and stability, proper mechanosensation and neuromuscular function. Notably, our analyses suggest that NDG could be an important factor to ensure ECM plasticity in response to environmental cues.

## MATERIALS AND METHODS

### Fluorescence antibody staining

Antibody staining of *Drosophila* embryos and larvae was essentially performed as described by Mueller in (Dahmann, 2008). For antibody staining of wandering 3^rd^ instar larvae we adapted the protocol from Klein in (Dahmann, 2008) with the following modifications: animals were relaxed before dissection by briefly dipping them into 60 °C water and a permeabilization step (10 min wash in PBS supplied with 1% Triton-X100) was added before blocking and primary antibody incubation. The following primary antibodies were used in their specified dilutions: mouse anti-Bruchpilot (Brp nc82, 1:100, (Wagh et al., 2006), DSHB), guinea pig anti-Collagen IV (COLLIV, 1:500, (Lunstrum et al., 1988)), sheep anti-Digoxygenin alkaline phosphatase conjugated Fab fragments (DIG-AP, 1:4.000, Roche Applied Science), mouse anti-Green fluorescent protein (GFP, 1:500, Roche Diagnostics), rabbit anti-Green fluorescent protein (GFP, 1:500, abcam), chicken anti-Green fluorescent protein (GFP, 1:500, abcam), goat anti-Horseradish peroxidase, Cy3-conjugated (HRP, 1:200, Jackson ImmunoResearch), guinea pig anti-Laminin A (LanA, 1:500, (Harpaz and Volk, 2012)), mouse anti-Laminin A (LanA, 1:500, (Takagi et al., 1996)), rabbit anti-Laminin B1 (LanB1, 1:400, (Kumagai et al., 1997), abcam), rabbit anti-Laminin B2 (LanB2, 1:400, (Kumagai et al., 1997), abcam), rabbit anti-K-cell Laminin (LanKc, 1:500, (Gutzeit et al., 1991)), rabbit anti-Wing blister (LanWb, 1:100, (Martin et al., 1999)), rabbit anti-Nidogen (NDG, 1:1.000, (Wolfstetter et al., 2009)), rabbit anti-Perlecan (PCAN, 1:1.000, (Friedrich et al., 2000)) and rabbit anti-SPARC (1:500, (Martinek et al., 2008)). Alexa Fluor-, Cy-, Biotin-SP-, and HRP-coupled secondary antibodies were purchased from Dianova and Jackson ImmunoResearch, DAPI from Sigma Aldrich. Embryos and larval tissues were embedded in Fluoromount-G (Southern Biotech) before visualization under Leica TCS SP2 or Zeiss LSM 800 confocal microscopes.

### Developmental Northern blot

Northern blot analyses were performed by standard procedures (Maniatis, 1982). RNA was extracted by the guanidine thiocyanate/phenol/chloroform extraction method of Chomczynski and Sacchi (Chomczynski and Sacchi, 1987). Poly(A)-tailed RNA was isolated using a Pharmacia Kit (Pharmacia Biotech). A 2.5 kb fragment of the *Ndg* cDNA *(cNdg5*, Stefan Baumgartner unpublished) was radioactively labelled and hybridized to the Northern filter. Exposure time for Northern blots was 1.5 d. The blot was re-probed with a *Drosophila 19S* probe to evaluate equimolar loading. The *Ndg* blot shown in this work was identically conducted as those for *wing blister (LanWb)* and *LanA* (Martin et al., 1999) allowing scarce comparison of the expression levels of these different ECM genes.

### Whole mount *in situ* hybridization

N-and C-terminal fragments from a full-length *Nidogen* cDNA clone (DGRC cDNA clone LP19846; GenBank: BT031149.1) were PCR-amplified and sub-cloned into the pCRII-TOPO vector with the TOPO TA Dual Promoter Kit (Invitrogen). Primer sequences for a 571 bp N-terminal fragment were GGACCCATCCATATCCCGCCACAAT and GCAATCAGTGCCACCTGGAAGGTGT while CGTGGCATTGCCGTGGATCCCT and GGTGCATCCTGTGGAGGCGCT were employed to amplify a 549 bp C-terminal fragment. Templates for probe synthesis were generated by PCR using M13 primers supplied with the TOPO TA Dual Promoter Kit. Digoxygenin (DIG)-labeled *sense* and *antisense* probes were made by SP6/T7 *in vitro* transcription with the DIG RNA Labeling Kit (Roche Applied Science). *In situ* hybridization on *Drosophila* embryos was performed according to (Lecuyer et al., 2008) with modifications adapted from (Pfeifer et al., 2012). Sheep anti-DIG-AP Fab fragments (1:4.000, Roche Applied Science), biotinylated donkey anti-sheep IgG (1:400, Dianova), the Vectastain ABC Standard Kit (Vector Laboratories) and the TSA Amplification Renaissance Kit (PerkinElmer) were used for fluorescence *in situ* hybridization (FISH) detection. Specificity of the probes was tested by *in situ* hybridization on *white^1118^*, and *Ndg* deficient *Df(2R)BSC281* embryos, whereupon we detected no differences between the N-and the C-terminal probes (not shown).

### Fly stocks and genetics

Flies were grown under standard conditions (Ashburner, 1989) and crosses were performed at room temperature or at 25 °C. Staging of embryos was done according to (Campos-Ortega and Hartenstein, 1997). The following mutations and fly stocks were used in this study: as control or wild-type stocks we employed *white*^*1118*^ (*w*^*1118*^), *w*; Kr^lf-1^/CyO P{Dfd-EYFP}2, Oregon-R* or balanced sibling embryos. We used *Df(2R)BSC281* as a deficiency for *Ndg (Df Ndg), Df(2L)Exel7022* which deletes the two adjacent *Drosophila* type IV Collagen genes *viking (vkg)* and *Col4a1* (previously referred to as *Cg25C), Df(3L)Exel8101* as deficiency for *Laminin A, Df(2L)TE35B-2* as deficiency for *Laminin wing blister* (Gubb et al., 1985), *Df(3L)Exel6114* as *Laminin B2* deficiency *(Df LanB2)* and *Df(1)Exel6230* that removes the *trol* locus but also the adjacent segmentation gene *giant. Mi{ET1}MB04184, P{hsILMiT}* and *P(A2-3}99B* were used to generate the *Ndg* deletion alleles *Ndg^A04^.* and *Ndg*^*Δ1.4*^. (Sup. Fig 2, see below). The null alleles *LanB2^knod^.* (Wolfstetter and Holz, 2012) and *wb^HG10^.* were used as well as *Lar*^13.2^ that encodes for a truncated form of the Leukocyte-antigen-related-like receptor (Krueger et al., 1996). Protein trap lines were: *vkg::GFP* (Morin et al., 2001), *trol::GFP* (Medioni and Noselli, 2005), and *Jupiter::GFP* (Karpova et al., 2006).

### Generation of *Nidogen* deletions

The *pMiET1* transposon insertion *Mi{ET1}MB04184* (Metaxakis et al., 2005) was used in an imprecise excision screen. Lethality reported for this line was not associated with the *Mi{ET1}* insertion and homozygous viable, isogenic stocks were established after ~5 generations of free recombination over a wild-type chromosome. The position of the *Mi{ET1}* insertion, initially revealed by flanking sequence recovery ((Bellen et al., 2011), GenBank: ET201740.1), was confirmed in the isogenic stocks by inverse PCR (Ochman et al., 1988). Thereby, we detected a 35 bp deletion 141 bp upstream of the insertion site that was present in the flanking sequence recovery data (ET201740.1) and in *w*^*1118*^. DNA samples but not in the 6^th^ GenBank release of the *,D. melanogaster*, genome annotation (Hoskins et al., 2015) or the corresponding sequence derived from *Oregon-R* genomic DNA. Therefore, we considered this small deletion as a naturally occurring sequence variation in the *Ndg* locus. Before serving as Minos transposase source in the screen, the *P{hsILMiT}* insertion (Metaxakis et al., 2005) was remobilized from the *SM6a* balancer and inserted onto a *w*^*1118*^. X-chromosome employing *P(A2-3}99B* as transposase source. *pMiET1* remobilization was induced in the germ line of 2^nd^ and 3^rd^ instar larvae by daily 1 h heat shocks at 37 °C in the presence of the *w*^*1118*^, *P{hsILMiT}* helper chromosome. A total of 301 single excision events were identified due to the absence of the *Mi{ET1}-associated Mmus/Pax6-GFP* expression (Berghammer et al., 1999) and screened for deletions by single fly PCR analysis with the following primer combination: G C C AAG G AAT GGGAGTGCTCTGGAT and GGAGCCATCCTCGAACTCGTACAATT. Deletions were detected in two viable *Mi{ET1}MB04184* excision lines (henceforth referred to as *Ndg*^*Δ0.4*^. and *Ndg*^*Δ1.4*^). Sequencing the molecular lesions uncovered a 1401 bp deletion (2R: 10312780-10314180) and a P-Element remnant of 17 nucleotides (TGCCACGTAGCCGGAAT) in *Ndg^*Δ1.4*^* In the case of *Ndg*^*Δ0.4*^. we detected a 417 bp (2R:10313549-10313965) deletion and an insertion of 19 nucleotides (CGAGCAAAATACAAAATAC) at the former P-Element insertion site (Sup. Fig 2).

### Dextran permeability assay

Wandering 3^rd^ instar larvae were picked and relaxed by briefly dipping them into 60 °C water. Larval filets were dissected and incubated in 50 Ml of 25 Mg/ml anionic Texas Red-coupled Dextran (10.000 MW, Thermo Fischer Scientific) in PBS for 10 minutes. The dextran solution was removed and the larval filets were rinsed once with 50 μl PBS. Larval filets were fixed for 10 min in a drop of 4% formaldehyde in PBS and rinsed with PBS before mounting in Fluoromount-G and analysis under a Zeiss LSM 800 confocal microscope.

### Larval climbing and gravity assay

Matching numbers of flies were placed on food-filled vials and allowed to lay eggs for three days. Pupae were examined at the pharate adult stage (7-9 days after egg laying at 25 °C). To assay larval climbing performance, the distance between the food surface and the cotton plug was divided into four zones. Pupae formed in each zone were counted. Pupae positioned on the border between two zones were assigned to the next higher zone. Pupae were further assigned into three categories (1. Upright = 0° “head up” or 180° “head down” +/− 22.5°, 2. Tilted = 45° or 225° +/− 22.5°, and 3. Flat = 90° or 270° +/− 22.5°) according to their orientation along the gravity axis.

### Scanning electron microscopy (SEM)

Wandering 3^rd^ instar larvae were dissected in PBS. After 3 h fixation in 2.5 % glutaraldehyde in PBS, samples were rinsed several times in PBS and dehydrated in an ascending ethanol series (50 %, 70 %, 80 %, 90 %, and 2x 100 % ethanol for 10 min each). After critical-point-drying in a Balzers CPD 030 at 40 °C, samples were mounted on stubs using double-sided adhesive tape and sputter-coated with a thin layer of gold (Balzers SCD 004). Samples were analyzed under a Zeiss DSM982 scanning electron microscope. Acceleration voltage was set to 3 kV. Settings for tilt angle, spot size, scanning mode and magnifications were kept constant throughout image acquisition.

### Osmotic stress assay

Wandering 3^rd^ instar larvae of indicated genotypes were picked and rinsed with PBS. The larvae were cut in half and the anterior part was inverted in order to expose the wing imaginal discs. After removing the anterior part of the gut and fat body, the samples were transferred to distilled water and the time until bursting of the first wing imaginal disc was measured. Measurements were stopped after 10 min and wing imaginal discs that were found intact after this period were assigned to the 10 min group.

### Lethality tests

Parental flies (3-7 days old) of the indicated genotype were shifted on grape juice agar plates for 8 days. Agar plates were changed every 2^nd^ day and defined numbers of eggs were transferred to fresh agar plates supplemented with dried yeast for further comparative analysis. Undeveloped eggs, hatched larvae, pupal cases and enclosure were counted daily. All plates were incubated at 25 °C and constantly humidified.

### Larval feeding assay

Fifty freshly hatched 1^st^ instar larvae were transferred to apple juice agar plates with a central spot of food-dye-supplemented yeast paste. Larvae moving outside the food were counted 1 and 2 h after the transfer. Fresh yeast paste was added every day and individual larvae were checked for the uptake of the dyed food after 72 h. The assay was performed at 25 °C and 60 % humidity conditions.

### Larval locomotion assay

Flies were allowed to lay eggs for 2 h on apple juice agar plates supplemented with yeast paste. Fresh yeast paste was added every day and individual 2^nd^ instar larvae were picked after 72 h (at 25 °C and 60 % humidity) and rinsed with distilled water. A single larva was placed in the middle of a 0 52.5 mm Petri dish prepared with a 0 35 mm central arena (1 % agar in PBS) surrounded by a ring of high salt agar (3 M NaCl in PBS) as locomotion barrier. For every recording a fresh agar plate was used. Larvae were allowed to move freely in the central arena for 2-5 min respectively and locomotion was recorded in a square of 12,5×9,5 mm^2^ using a Zeiss AxioZoom.V16 stereo zoom microscope equipped with LED ring light and an Axiocam 503 color camera. Time series were recorded with 4 frames/second employing ZEN Blue software and 25 s subsets were further analyzed and edited with the manual tracking plugin MTrackJ and the Fiji distribution of ImageJ (Schindelin et al., 2012).

### Larval crawling assay

Larval crawling analyses were performed as described by (Eschbach et al., 2011) for responses to vibrational stimulation and measurement of body length and stride frequency. For the trials, 3^rd^ instar larvae were placed on 1 % agarose covered Petri dishes (60 mm diameter) as a crawling stage. This stage was placed in a silencing foam covered box (57×57×47 cm^3^.) and illuminated by a red darkroom lamp (Pf712em, Philips, 15 W and 7 lm). Video recordings of the trials were performed via an infrared CMOS-Color-Camera (CCD-651, Conrad Electronics). The analogue video signals were grabbed by using an analogue/digital video converter (700-USB, Pinnacle Systems) and recorded via a computer for further analysis (25 fps). Measurements of crawling speed, stride frequency and body length of the larvae were performed by using the video tracking software Cabrillo Tracker (v4.91, Douglas Brown, Open Source Physics, 2015). To quantify mean velocity and stride frequency for five larvae per trial, the terminal ends were video-tracked for 10-20 s. Only periods of unrestricted and constant crawling behavior were used for analysis. The mean velocity was calculated from 5 s of crawling and the stride frequency from the duration of 5 complete strides in a row. The maximum crawling speed of a single stride was defined as the starting point for each measurement. To determine body length reduction due to vibrational stimuli, apical and terminal ends of four larvae per trial were video-tracked. The vibrational stimuli were generated via the audio edition software Audacity (v2.05, Audacity-Team, GNU GPLv2+, 2015) at a frequency of 100 Hz and delivered via a full-range speaker (BPSL 100/7, Isophon/Gauder Akustik, 60-20.000 Hz and 7 W) to the crawling stage. For each trial, three stimuli with the duration of 1 s and an acceleration of 12 m/s^2^ were applied (interpulse-interval 9 s).

### Statistical analyses

Prism v6.0 and v7.02 (GraphPad Software Inc.) was used for all statistical analyses.

## Acknowledgements

Bloomington *Drosophila* Stock Center (NIH P40OD018537), Lynn Cooley, the *Drosophila* Genomics Resource Center (DGRC, NIH 2P40OD010949-10A1), the Developmental Studies Hybridoma Bank (created by the NICHD of the NIH and maintained at the University of Iowa), Liselotte Fessler, the KYOTO Stock Center at the Kyoto Institute of Technology, Maurice Ringuette, John Roote, Talila Volk and Gerd Vorbrüggen for sending materials and fly stocks.

## Funding

This work was supported by:

SB: Vetenskapsrådet (621-2003-3408) and Cancerfonden (4714-B03-02XBB).

RHP: Vetenskapsrådet (621-2015-04466), Cancerfonden (2015/391), Barncancerfonden (2015/0096), and Göran Gustafsson Stiftelser.

AH: Deutsche Forschungsgemeinschaft/DFG (Ho-2559/3-3 and 2559/5-1).

## Author contributions

**GW:** Investigation, methodology, conceptualization, visualization, data curation, formal analysis, supervision, validation, writing original draft, writing review and editing.

**ID:** Investigation, visualization, formal analysis, validation and writing original draft.

**KP:** Investigation, visualization, formal analysis, validation and writing original draft.

**JA:** Investigation, visualization, formal analysis, validation and writing original draft.

**UT**: Investigation, visualization, formal analysis, validation and writing original draft.

**DP:** Investigation, visualization and writing original draft.

**RL-H:** Supervision.

**SB:** Investigation, writing original draft and funding acquisition, writing review and editing.

**RHP:** Supervision, funding acquisition, writing review and editing.

**AH:** Investigation, conceptualization, methodology, visualization, supervision, validation, funding acquisition, project administration, writing original draft, writing review and editing.

## Supplementary Figure Legends

**Supplementary Figure 1:**
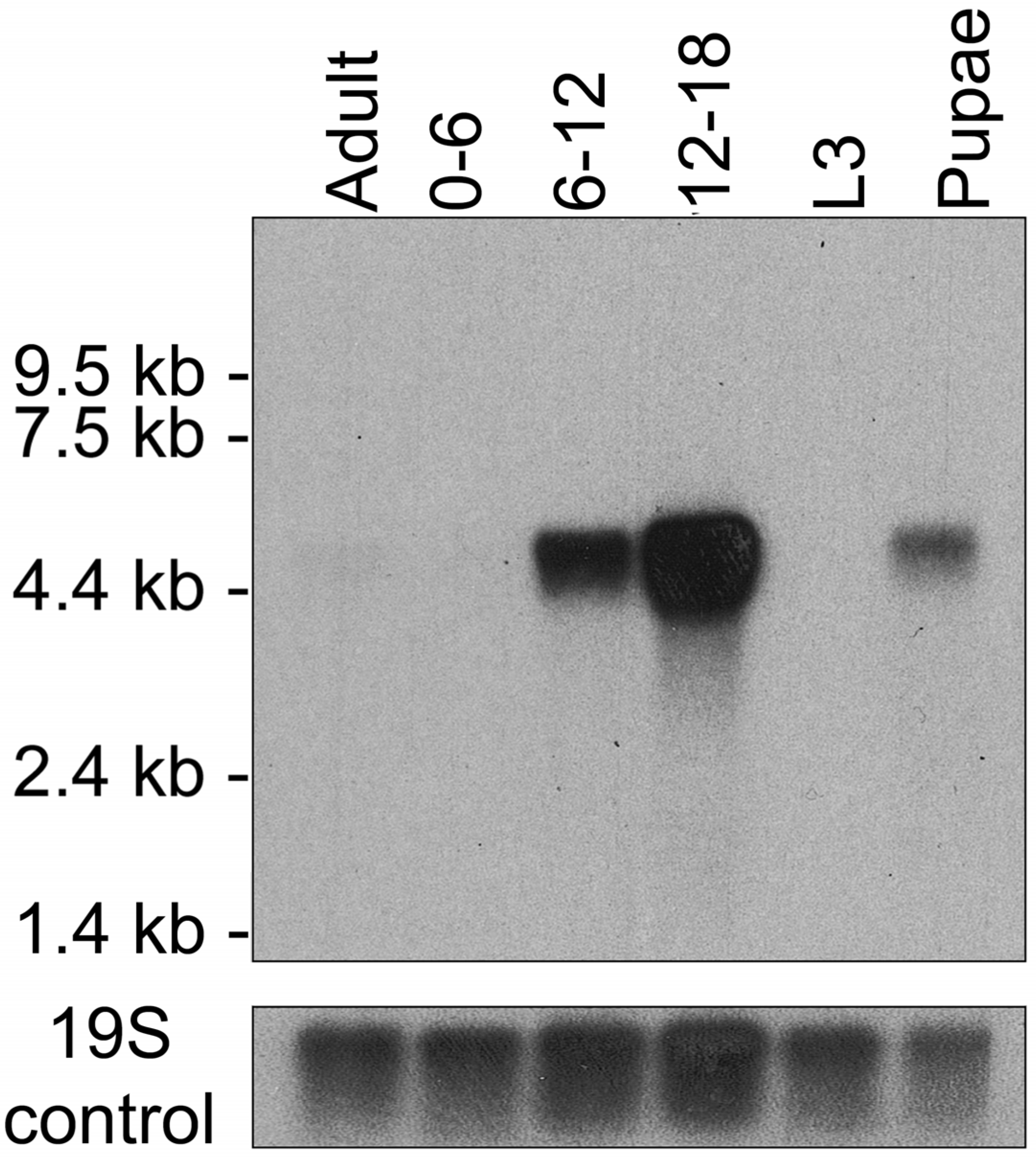
Developmental Northern blot analysis of *Ndg* expression. Total *mRNA* from adult flies (lane 1), embryos from 0-6 h, 6-12 h, 12-18 h (lanes 2-4), 3^rd^ instar larvae (L3, lane 5) and pupae was extracted and detected by an Ndg-specific probe. No signal was detectable up to 6 h of embryonic development as well as in L3 larvae and adult flies. Later embryonic stages and pupae exhibit *Ndg mRNA* expression with the strongest intensity around 12 to 18 h of embryonic development. 19S rRNA was used as loading control.>

**Supplementary Figure 2:**
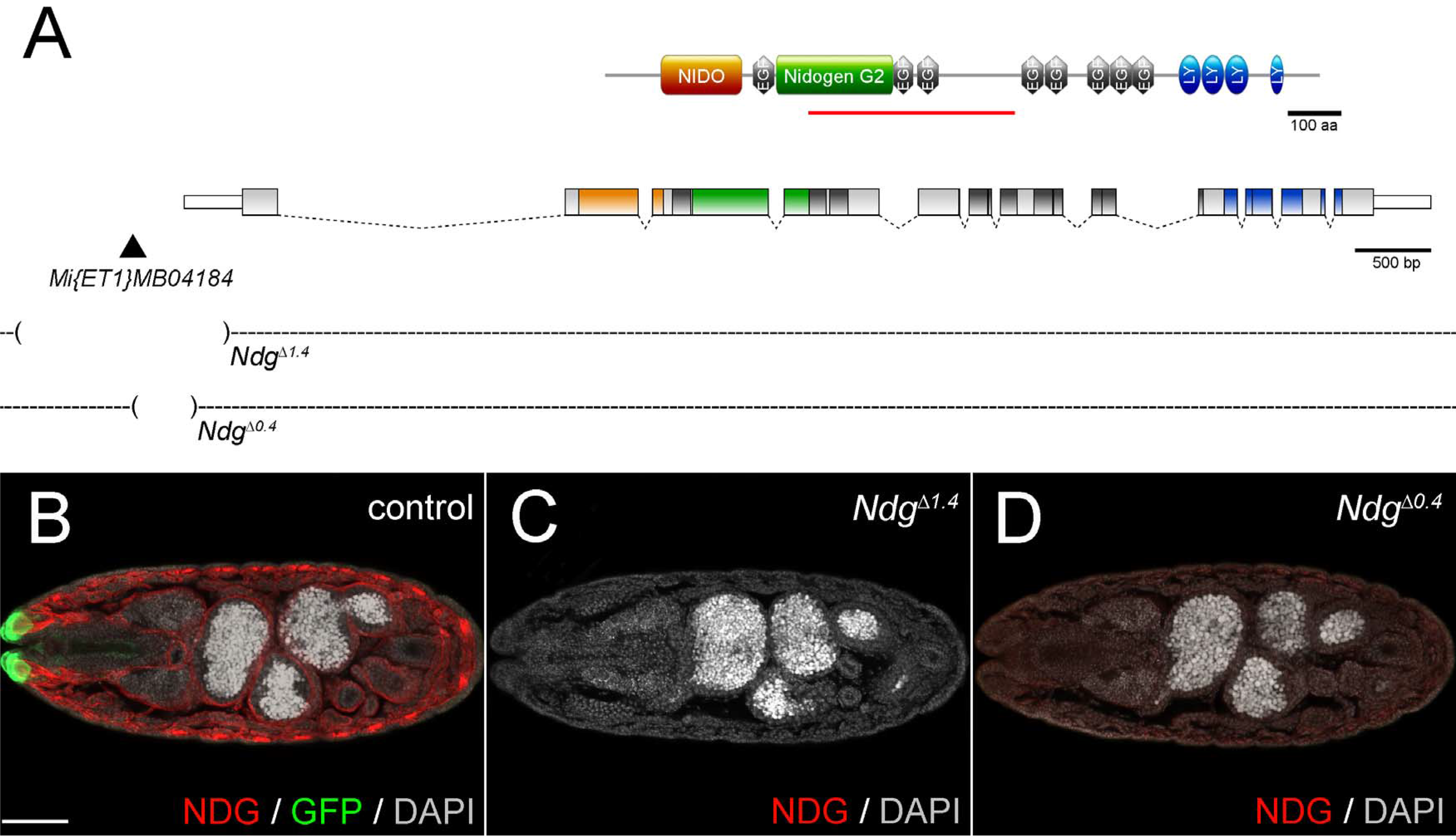
Generation of *Ndg* mutants. (**A**) Schematic representation (created with (Hulo et al., 2008;Rambaldi and Ciccarelli, 2009)) of the NDG protein domain organization (**top**) and the *Ndg* locus (**below**) including the positions of the mobilized P-element *Mi{ET1}MB04184* and the obtained genomic deletions *Ndg^A1′^* ^4^ and *Ndg^Δ0.4;^* ^4^. Protein domains and parts of the locus that encode for these domains are highlighted in matching colors. The position of the epitope employed to generate the NDG antibody (Wolfstetter et al., 2009) is indicated by a red line. NIDO: NIDO domain, EGF: EGF-like domain, Nidogen G2: Nidogen G2 p-barrel domain, LY:.LDL-receptor class B (**LDLRB**)/YWTD repeat. (**B-D**) Confocal stacks from dorsal views of stage 16 *Drosophila* embryos stained for NDG (**red**). GFP (**green**) depicts balancer-associated reporter gene expression in a sibling control. DAPI and auto fluorescence of yolk droplets appears in one channel (**white**). (**B**) Strong NDG staining of BMs in control embryos. (**C**) Absence of detectable NDG protein in embryos homozygous for the *Ndg*^*Δ1.4*^. deletion. (**D**) Decreased NDG protein levels in a homozygous *Ndg*^*Δ0.4*^. embryo. Confocal images were acquired with identical settings for the Cy3-detecting channel. Scale bar = 50 μm.

**Supplementary Figure 3:**
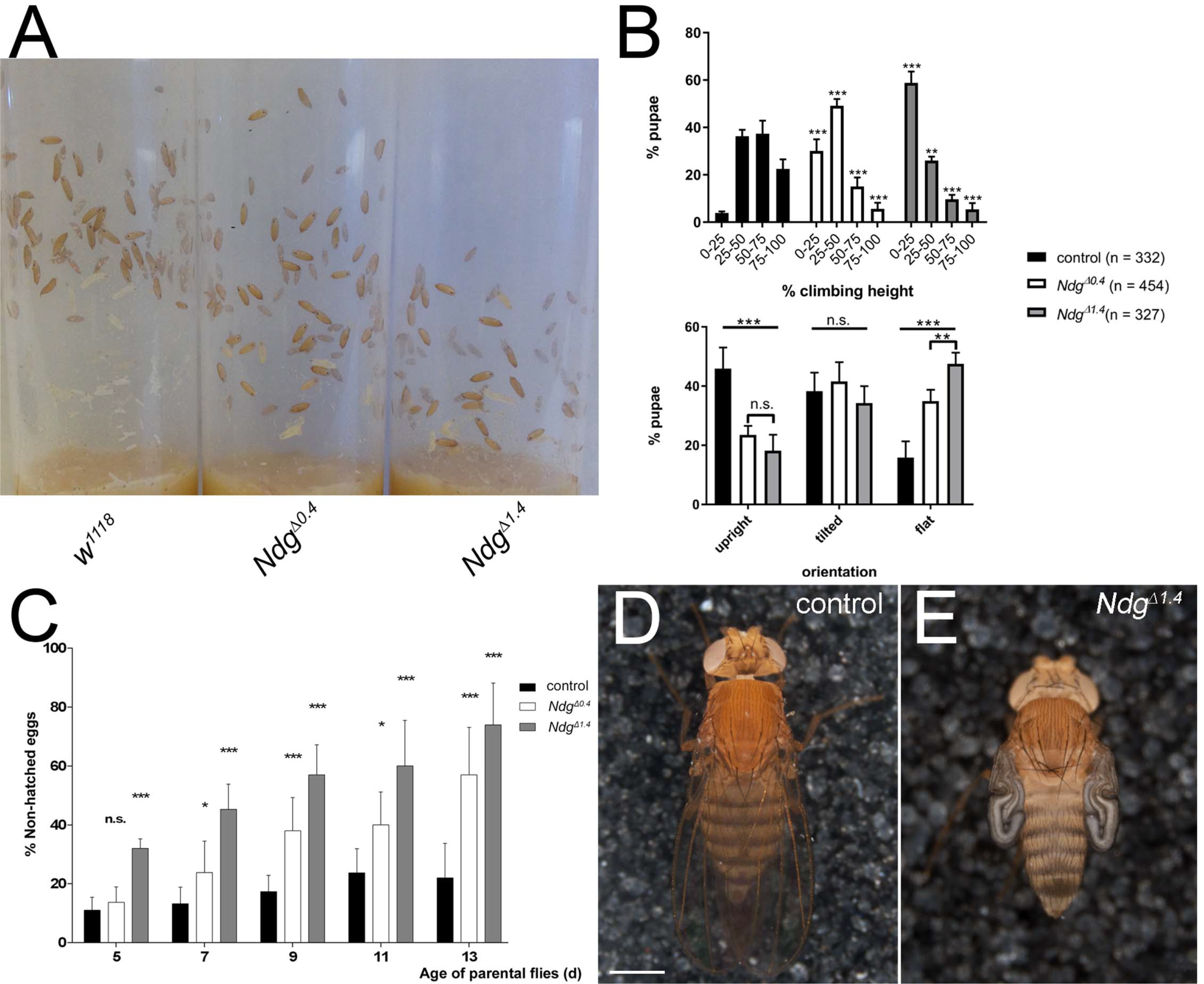
*Ndg* mutant larvae exhibit motility and gravity-sensing defects whereas *Ndg* mutant flies display decreased fecundity. (**A**) Formation of pupal cases in vials of *white^1110^* control and homozygous Ndg^Δ^. stocks (matching numbers of parental flies and breeding days at 25 °C). (**B**) Quantification of the climbing and orientation phenotypes. In comparison to the control, *Ndg*^*Δ*^ pupal cases are preferentially found in the lower half of the vials indicating impaired climbing performance (2-W anova and Tuckey’s multiple comparisons test with adjusted p-values, *** = p < 0.001, **p = 0.005, non-significant (n.s.) = p > 0.05). In *Ndg^Δ^* vials, the orientation of the pupal cases is significantly shifted towards the horizontal axis (2-W anova and Tuckey’s multiple comparisons test with adjusted p-values, ***p < 0.001, **p = 0.008, non-significant (n.s.) = p > 0.05). (**C**) Percentage of non-hatched eggs produced by 5 to 13 days old control and *Ndg*^*Δ*^ flies incubated at 25 °C. *Ndg*^*Δ*^ produce a significantly increasing number of non-hatched eggs over time, (2-W anova and Tuckey’s multiple comparisons test with adjusted p-values, *** = p < 0.001, ** = p < 0.01, * = p < 0.05 non-significant (n.s.) = p > 0.05). (**D**) Image of a control fly exhibiting properly inflated wings (n = 3492 animals). (**E**) Impaired wing inflation is observed at low penetrance in *Ndg*^*Δ1.4*^. flies (140 of n = 2839 animals). Scale bar = 500 μm.

**Supplementary Figure 4:**
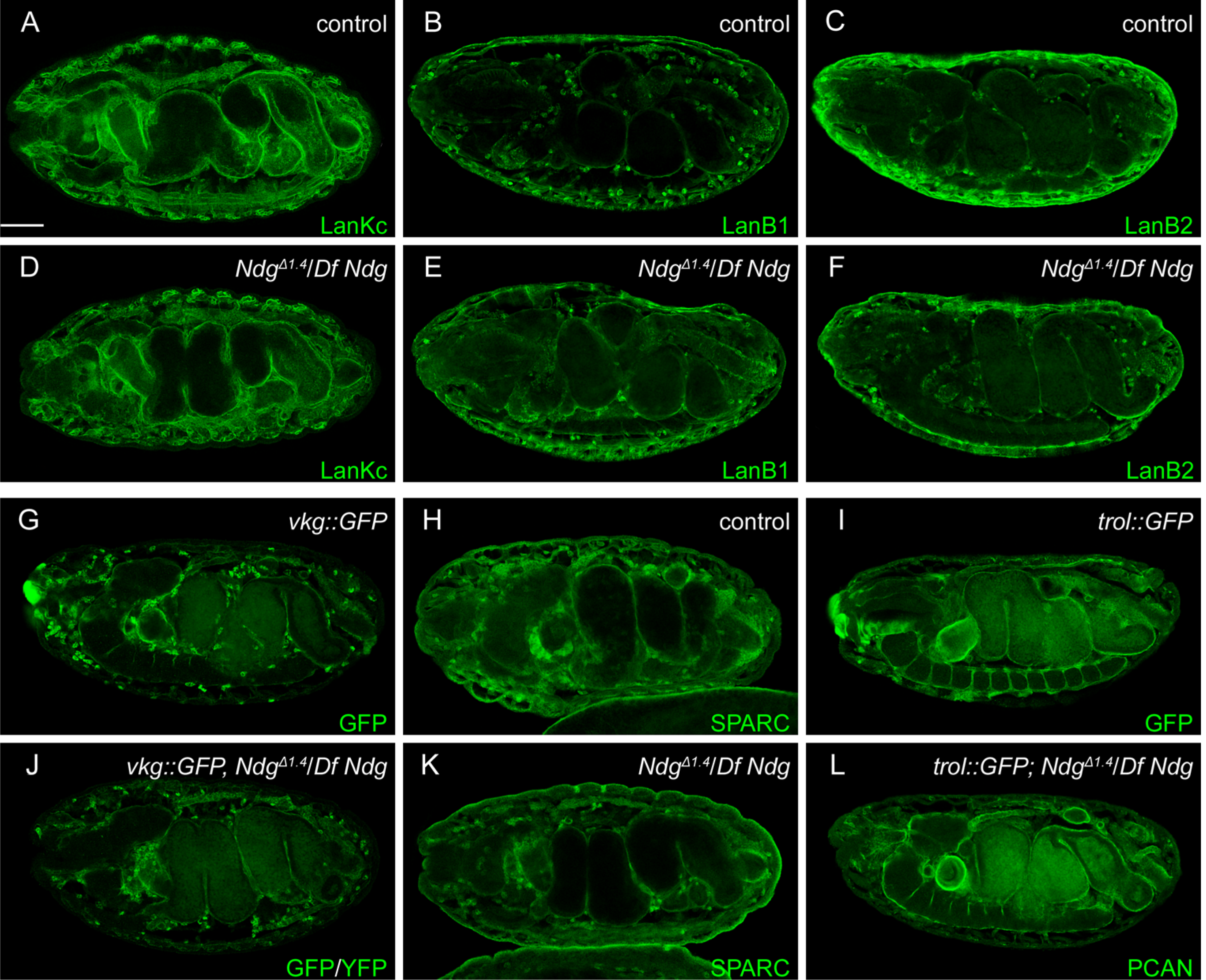
BM formation in the absence of *Ndg.* (A, B, C) Antibody staining against LanKc, LanB1 or LanB2 labels BMs, and in the case of LanB1 and LanB2 additionally fat body and hemocytes of control and transheterozygous *Ndg^*Δ1.4*^IDf Ndg* (**D**, **E**, **F**) embryos at stage 16. (**G**, **H**, **J**, **K**) A similar localization pattern was observed for *vkg::GFP* and SPARC in control and *Ndg^*Δ1.4*^IDf Ndg* embryos. (*I, L*) *trol::GFP* localization in control and *Ndg^AA4^/Df Ndg* embryos. Embryos appear in lateral (**B, C, E, F, G, I, K, L**) or ventrolateral (**A**, **D**, **H**, **J**) orientation. Scale bar = 50 μm.

**Supplementary Figure 5:**
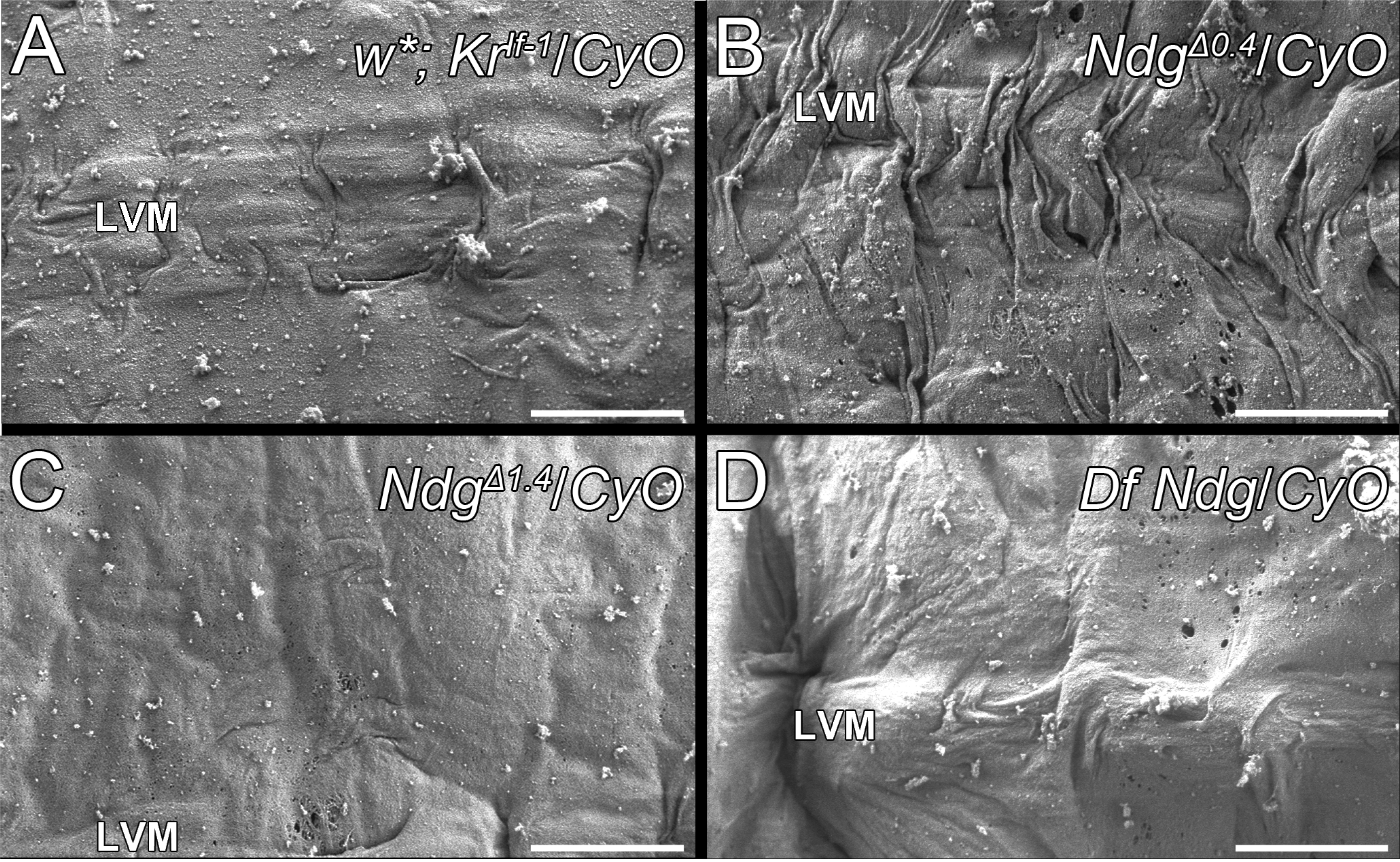
Heterozygous *Ndg*^*Δ*^ animals display a weak BM phenotype. (A-D) BM ultrastructure of (**A**) w*; *Kμ/CyO* control 3^rd^. instar larvae and (B-D) heterozygous *Ndg* mutants with indicated genotype revealed by SEM. LVM: Longitudinal visceral muscle. Scale bars = 5 μm.

**Supplementary Figure 6:**
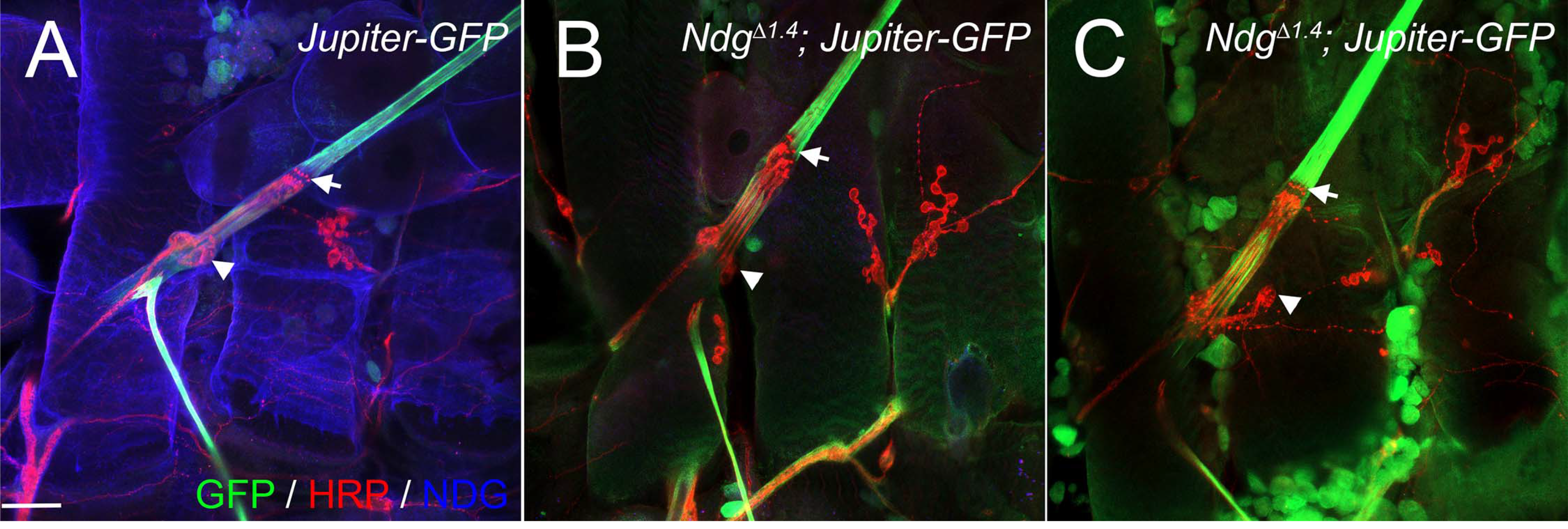
Chordotonal organs display aberrant neuron morphology in the absence of NDG. (**A**) Jupiter::GFP fusion protein (**green**) localizes to the microtubule network of the chordotonal organ in a *Jupiter-GFP* control larva. The chordotonal organ is surrounded by a NDG containing BM (**blue**) that also covers HRP-labelled neurons (red, arrowhead) and sensory cilia (**arrows**). (B, C) Two hemisegments of a *Jupiter-GFP; Ndg^A14^.* larva exhibit neuron detachment (**arrowheads**) and alignment defects of sensory cilia (**arrow**) in the lateral pentascolopidial chordotonal organ. Scale bar = 50 μm.

**Supplementary Movie:** Recordings of crawling patterns from *white^1118^* control and *Ndg^A14^.* 1st instar larvae.

## References

Ackley, B. D., Harrington, R. J., Hudson, M. L., Williams, L., Kenyon, C. J., Chisholm, A. D. and Jin, Y. (2005). The two isoforms of the Caenorhabditis elegans leukocyte-common antigen related receptor tyrosine phosphatase PTP-3 function independently in axon guidance and synapse formation. J Neurosci 25, 7517–7528.

Ackley, B. D., Kang, S. H., Crew, J. R., Suh, C., Jin, Y. and Kramer, J. M. (2003). The basement membrane components nidogen and type XVIII collagen regulate organization of neuromuscular junctions in Caenorhabditis elegans. J Neurosci 23, 3577–3587.

Arikawa-Hirasawa, E., Watanabe, H., Takami, H., Hassell, J. R. and Yamada, Y. (1999). Perlecan is essential for cartilage and cephalic development. Nat Genet 23, 354–358.

Artero, R., Furlong, E. E., Beckett, K., Scott, M. P. and Baylies, M. (2003). Notch and Ras signaling pathway effector genes expressed in fusion competent and founder cells during Drosophila myogenesis. Development 130, 6257–6272.

Ashburner, M. (1989). Drosophila: A laboratory manual: Cold Spring Harbor Laboratory.

Aumailley, M., Battaglia, C., Mayer, U., Reinhardt, D., Nischt, R., Timpl, R. and Fox, J. W. (1993). Nidogen mediates the formation of ternary complexes of basement membrane components. Kidney Int 43, 7–12.

Bader, B. L., Smyth, N., Nedbal, S., Miosge, N., Baranowsky, A., Mokkapati, S.,Murshed, M. and Nischt, R. (2005). Compound genetic ablation of nidogen 1 and 2 causes basement membrane defects and perinatal lethality in mice. Mol Cell Biol 25, 6846–6856.

Bellen, H. J., Levis, R. W., He, Y., Carlson, J. W., Evans-Holm, M., Bae, E., Kim, J., Metaxakis, A., Savakis, C., Schulze, K. L., et al. (2011). The Drosophila gene disruption project: progress using transposons with distinctive site specificities. Genetics 188, 731–743.

Berghammer, A. J., Klingler, M. and Wimmer, E. A. (1999). A universal marker for transgenic insects. Nature 402, 370–371.

Bharadwaj, R., Roy, M., Ohyama, T., Sivan-Loukianova, E., Delannoy, M., Lloyd, T. E., Zlatic, M., Eberl, D. F. and Kolodkin, A. L. (2013). Cbl-associated protein regulates assembly and function of two tension-sensing structures in Drosophila. Development 140, 627–638.

Borchiellini, C., Coulon, J. and Le Parco, Y. (1996). The function of type IV collagen during Drosophila embryogenesis. Roux Arch Dev Biol 205, 468–475.

Bose, K., Nischt, R., Page, A., Bader, B. L., Paulsson, M. and Smyth, N. (2006). Loss of nidogen-1 and −2 results in syndactyly and changes in limb development. J Biol Chem 281, 39620–39629.

Busser, B. W., Gisselbrecht, S. S., Shokri, L., Tansey, T. R., Gamble, C. E., Bulyk, M. L. and Michelson, A. M. (2013). Contribution of distinct homeodomain DNA binding specificities to Drosophila embryonic mesodermal cell-specific gene expression programs. PLoS One 8, e69385.

Busser, B. W., Taher, L., Kim, Y., Tansey, T., Bloom, M. J., Ovcharenko, I. and Michelson, A. M. (2012). A machine learning approach for identifying novel cell type-specific transcriptional regulators of myogenesis. PLoS Genet 8, e1002531.

Campos-Ortega, J. A. and Hartenstein, V. (1997). The embryonic development of Drosophila melanogaster. 2nd ed.

Carlin, B., Jaffe, R., Bender, B. and Chung, A. E. (1981). Entactin, a novel basal lamina-associated sulfated glycoprotein. J Biol Chem 256, 5209–5214.

Chiang, C., Patel, N. H., Young, K. E. and Beachy, P. A. (1994). The novel homeodomain gene buttonless specifies differentiation and axonal guidance functions of Drosophila dorsal median cells. Development 120, 3581–3593.

Chomczynski, P. and Sacchi, N. (1987). Single-step method of RNA isolation by acid guanidinium thiocyanate-phenol-chloroform extraction. Anal Biochem 162, 156–159.

Clay, M. R. and Sherwood, D. R. (2015). Basement Membranes in the Worm: A Dynamic Scaffolding that Instructs Cellular Behaviors and Shapes Tissues. Curr Top Membr 76, 337–371.

Dahmann, C. (2008). Drosophila: Methods and Protocols: Humana Press.

Datta, S. and Kankel, D. R. (1992). l(1)trol and l(1)devl, loci affecting the development of the adult central nervous system in Drosophila melanogaster. Genetics 130, 523–537.

Dong, L., Chen, Y., Lewis, M., Hsieh, J. C., Reing, J., Chaillet, J. R., Howell, C. Y.,Melhem, M., Inoue, S., Kuszak, J. R., et al. (2002). Neurologic defects and selective disruption of basement membranes in mice lacking entactin-1/nidogen-. Lab Invest 82, 1617–1630.

Durkin, M. E., Bartos, B. B., Liu, S. H., Phillips, S. L. and Chung, A. E. (1988). Primary structure of the mouse laminin B2 chain and comparison with laminin B1. Biochemistry 27, 5198–5204.

Eschbach, C., Cano, C., Haberkern, H., Schraut, K., Guan, C., Triphan, T. and Gerber, B. (2011). Associative learning between odorants and mechanosensory punishment in larval Drosophila. J Exp Biol 214, 3897–3905.

Fox, J. W., Mayer, U., Nischt, R., Aumailley, M., Reinhardt, D., Wiedemann, H., Mann, K., Timpl, R., Krieg, T., Engel, J., et al. (1991). Recombinant nidogen consists of three globular domains and mediates binding of laminin to collagen type IV. EMBO J 10, 3137–3146.

Fox, M. A., Ho, M. S., Smyth, N. and Sanes, J. R. (2008). A synaptic nidogen: developmental regulation and role of nidogen-2 at the neuromuscular junction. Neural Dev 3, 24.

Friedrich, M. V., Schneider, M., Timpl, R. and Baumgartner, S. (2000). Perlecan domain V of Drosophila melanogaster. Sequence, recombinant analysis and tissue expression. Eur J Biochem 267, 3149–3159.

Gubb, D., Roote, J., Harrington, G., McGill, S., Durrant, B., Shelton, M. and Ashburner, M. (1985). A preliminary genetic analysis of TE146, a very large transposing element of Drosophila melanogaster. Chromosoma 92, 116–123.

Gutzeit, H. O., Eberhardt, W. and Gratwohl, E. (1991). Laminin and basement membrane-associated microfilaments in wild-type and mutant Drosophila ovarian follicles. J Cell Sci 100 (Pt 4), 781–788.

Halfter, W., Dong, S., Yip, Y. P., Willem, M. and Mayer, U. (2002). A critical function of the pial basement membrane in cortical histogenesis. J Neurosci 22, 6029–6040.

Harpaz, N. and Volk, T. (2012). A novel method for obtaining semi-thin cross sections of the Drosophila heart and their labeling with multiple antibodies. Methods 56, 63–68.

Hashimshony, T., Feder, M., Levin, M., Hall, B. K. and Yanai, I. (2015). Spatiotemporal transcriptomics reveals the evolutionary history of the endoderm germ layer. Nature 519, 219–222.

Henchcliffe, C., Garcia-Alonso, L., Tang, J. and Goodman, C. S. (1993). Genetic analysis of laminin A reveals diverse functions during morphogenesis in Drosophila. Development 118, 325–337.

Ho, M. S., Bose, K., Mokkapati, S., Nischt, R. and Smyth, N. (2008). Nidogens-Extracellular matrix linker molecules. Microsc Res Tech 71, 387–395.

Hobert, O. and Bulow, H. (2003). Development and maintenance of neuronal architecture at the ventral midline of C. elegans. Curr Opin Neurobiol 13, 70–78.

Hohenester, E. and Yurchenco, P. D. (2013). Laminins in basement membrane assembly. Cell Adh Migr 7, 56–63.

Hopf, M., Gohring, W., Mann, K. and Timpl, R. (2001). Mapping of binding sites for nidogens, fibulin-2, fibronectin and heparin to different IG modules of perlecan. J Mol Biol 311, 529–541.

Hoskins, R. A., Carlson, J. W., Wan, K. H., Park, S., Mendez, I., Galle, S. E., Booth, B.W., Pfeiffer, B. D., George, R. A., Svirskas, R., et al. (2015). The Release 6 reference sequence of the Drosophila melanogaster genome. Genome Res 25, 445–458.

Hulo, N., Bairoch, A., Bulliard, V., Cerutti, L., Cuche, B. A., de Castro, E., Lachaize, C., Langendijk-Genevaux, P. S. and Sigrist, C. J. (2008). The 20 years of PROSITE. Nucleic Acids Res 36, D245–249.

Hynes, R. O. (2012). The evolution of metazoan extracellular matrix. J Cell Biol 196, 671–679.

Jan, L. Y. and Jan, Y. N. (1982). Antibodies to horseradish peroxidase as specific neuronal markers in Drosophila and in grasshopper embryos. Proc Natl Acad Sci U S A 79, 2700–2704.

Jayadev, R. and Sherwood, D. R. (2017). Basement membranes. Curr Biol 27, R207–R211.

Kang, S. H. and Kramer, J. M. (2000). Nidogen is nonessential and not required for normal type IV collagen localization in Caenorhabditis elegans. Mol Biol Cell 11, 3911–3923.

Karpova, N., Bobinnec, Y., Fouix, S., Huitorel, P. and Debec, A. (2006). Jupiter, a new Drosophila protein associated with microtubules. Cell Motility and the Cytoskeleton 63, 301–312.

Kaufmann, N., DeProto, J., Ranjan, R., Wan, H. and Van Vactor, D. (2002). Drosophila liprin-alpha and the receptor phosphatase Dlar control synapse morphogenesis. Neuron 34, 27–38.

Kim, S. and Wadsworth, W. G. (2000). Positioning of longitudinal nerves in C. elegans by nidogen. Science 288, 150–154.

Kramer, J. M. (2005). Basement membranes. WormBook, 1–15.

Krueger, N. X., van Vactor, D., Wan, H. I., Gelbart, W. M., Goodman, C. S. and Saito, H.(1996). The transmembrane tyrosine phosphatase DLAR controls motor axon guidance in Drosophila. Cell 84, 611–622.

Kumagai, C., Kadowaki, T. and Kitagawa, Y. (1997). Disulfide-bonding between Drosophila laminin beta and gamma chains is essential for alpha chain to form alpha betagamma trimer. FEBS Lett 412, 211–216.

Le Parco, Y., Cecchini, J. P., Knibiehler, B. and Mirre, C. (1986). Characterization and expression of collagen-like genes in Drosophila melanogaster. Biol Cell 56, 217–226.

Lecuyer, E., Parthasarathy, N. and Krause, H. M. (2008). Fluorescent in situ hybridization protocols in Drosophila embryos and tissues. Methods Mol Biol 420, 289–302.

Lindsley, D. L. and Zimm, G. G. (1992). GENES. In The Genome of Drosophila Melanogaster, pp. 1–803. San Diego: Academic Press.

Lunstrum, G. P., Bachinger, H. P., Fessler, L. I., Duncan, K. G., Nelson, R. E. and Fessler, J. H. (1988). Drosophila basement membrane procollagen IV. I. Protein characterization and distribution. J Biol Chem 263, 18318–18327.

Maniatis, T. (1982). Molecular cloning : a laboratory manual /T. Maniatis, E.F. Fritsch, J. Sambrook. Cold Spring Harbor, N.Y: Cold Spring Harbor Laboratory.

Mann, K., Deutzmann, R., Aumailley, M., Timpl, R., Raimondi, L., Yamada, Y., Pan, T. C., Conway, D. and Chu, M. L. (1989). Amino acid sequence of mouse nidogen, a multidomain basement membrane protein with binding activity for laminin, collagen IV and cells. EMBO J 8, 65–72.

Martin, D., Zusman, S., Li, X., Williams, E. L., Khare, N., DaRocha, S., Chiquet-Ehrismann, R. and Baumgartner, S. (1999). wing blister, a new Drosophila laminin alpha chain required for cell adhesion and migration during embryonic and imaginal development. J Cell Biol 145, 191–201.

Martin, G. R. and Timpl, R. (1987). Laminin and other basement membrane components. Annu Rev Cell Biol 3, 57–85.

Martinek, N., Shahab, J., Saathoff, M. and Ringuette, M. (2008). Haemocyte-derived SPARC is required for collagen-IV-dependent stability of basal laminae in Drosophila embryos. J Cell Sci 121, 1671–1680.

Matsubayashi, Y., Louani, A., Dragu, A., Sanchez-Sanchez, B. J., Serna-Morales, E., Yolland, L., Gyoergy, A., Vizcay, G., Fleck, R. A., Heddleston, J. M., et al. (2017). A Moving Source of Matrix Components Is Essential for De Novo Basement Membrane Formation. Curr Biol 27, 3526–3534 e3524.

Mayer, U., Nischt, R., Poschl, E., Mann, K., Fukuda, K., Gerl, M., Yamada, Y. and Timpl, R. (1993). A single EGF-like motif of laminin is responsible for high affinity nidogen binding. EMBO J 12, 1879–1885.

Mayer, U., Zimmermann, K., Mann, K., Reinhardt, D., Timpl, R. and Nischt, R. (1995). Binding properties and protease stability of recombinant human nidogen. EurJ Biochem 227, 681–686.

Medioni, C. and Noselli, S. (2005). Dynamics of the basement membrane in invasive epithelial clusters in Drosophila. Development 132, 3069–3077.

Metaxakis, A., Oehler, S., Klinakis, A. and Savakis, C. (2005). Minos as a genetic and genomic tool in Drosophila melanogaster. Genetics 171, 571–581.

Meyer, F. and Moussian, B. (2009). Drosophila multiplexin (Dmp) modulates motor axon pathfinding accuracy. Dev Growth Differ 51, 483–498.

Miosge, N., Kother, F., Heinemann, S., Kohfeldt, E., Herken, R. and Timpl, R. (2000). Ultrastructural colocalization of nidogen-1 and nidogen-2 with laminin-1 in murine kidney basement membranes. Histochem Cell Biol 113, 115–124.

Mirre, C., Le Parco, Y. and Knibiehler, B. (1992). Collagen IV is present in the developing CNS during Drosophila neurogenesis. J Neurosci Res 31, 146–155.

Montell, D. J. and Goodman, C. S. (1989). Drosophila laminin: sequence of B2 subunit and expression of all three subunits during embryogenesis. J Cell Biol 109, 2441–2453.

Morin, X., Daneman, R., Zavortink, M. and Chia, W. (2001). A protein trap strategy to detect GFP-tagged proteins expressed from their endogenous loci in Drosophila. Proceedings of the National Academy of Sciences of the United States of America 98, 15050–15055.

Murshed, M., Smyth, N., Miosge, N., Karolat, J., Krieg, T., Paulsson, M. and Nischt, R. (2000). The absence of nidogen 1 does not affect murine basement membrane formation. Mol Cell Biol 20, 7007–7012.

Nakae, H., Sugano, M., Ishimori, Y., Endo, T. and Obinata, T. (1993). Ascidian entactin/nidogen. Implication of evolution by shuffling two kinds of cysteine-rich motifs. Eur J Biochem 213, 11–19.

Ochman, H., Gerber, A. S. and Hartl, D. L. (1988). Genetic applications of an inverse polymerase chain reaction. Genetics 120, 621–623.

Ohyama, T., Jovanic, T., Denisov, G., Dang, T. C., Hoffmann, D., Kerr, R. A. and Zlatic, M. (2013). High-throughput analysis of stimulus-evoked behaviors in Drosophila larva reveals multiple modality-specific escape strategies. PLoS One 8, e71706.

Olson, P. F., Fessler, L. I., Nelson, R. E., Sterne, R. E., Campbell, A. G. and Fessler, J. H. (1990). Glutactin, a novel Drosophila basement membrane-related glycoprotein with sequence similarity to serine esterases. EMBO J 9, 1219–1227.

Pastor-Pareja, J. C. and Xu, T. (2011). Shaping cells and organs in Drosophila by opposing roles of fat body-secreted Collagen IV and perlecan. Dev Cell 21, 245–256.

Pfeifer, K., Dorresteijn, A. W. and Frobius, A. C. (2012). Activation of Hox genes during caudal regeneration of the polychaete annelid Platynereis dumerilii. Dev Genes Evol 222, 165–179.

Philippakis, A. A., Busser, B. W., Gisselbrecht, S. S., He, F. S., Estrada, B., Michelson, A. M. and Bulyk, M. L. (2006). Expression-guided in silico evaluation of candidate cis regulatory codes for Drosophila muscle founder cells. PLoS Comput Biol 2, e53.

Poschl, E., Schlotzer-Schrehardt, U., Brachvogel, B., Saito, K., Ninomiya, Y. and Mayer, U. (2004). Collagen IV is essential for basement membrane stability but dispensable for initiation of its assembly during early development. Development 131, 1619–1628.

Rambaldi, D. and Ciccarelli, F. D. (2009). FancyGene: dynamic visualization of gene structures and protein domain architectures on genomic loci. Bioinformatics 25, 2281–2282.

Reinhardt, D., Mann, K., Nischt, R., Fox, J. W., Chu, M. L., Krieg, T. and Timpl, R. (1993). Mapping of nidogen binding sites for collagen type IV, heparan sulfate proteoglycan, and zinc. J Biol Chem 268, 10881–10887.

Rodriguez, A., Zhou, Z., Tang, M. L., Meller, S., Chen, J., Bellen, H. and Kimbrell, D. A. (1996). Identification of immune system and response genes, and novel mutations causing melanotic tumor formation in Drosophila melanogaster. Genetics 143, 929–940.

Rossi, A., Kontarakis, Z., Gerri, C., Nolte, H., Holper, S., Kruger, M. and Stainier, D. Y. (2015). Genetic compensation induced by deleterious mutations but not gene knockdowns. Nature 524, 230–233.

Schindelin, J., Arganda-Carreras, I., Frise, E., Kaynig, V., Longair, M., Pietzsch, T., Preibisch, S., Rueden, C., Saalfeld, S., Schmid, B., et al. (2012). Fiji: an open-source platform for biological-image analysis. Nat Methods 9, 676–682.

Schymeinsky, J., Nedbal, S., Miosge, N., Poschl, E., Rao, C., Beier, D. R., Skarnes, W. C., Timpl, R. and Bader, B. L. (2002). Gene structure and functional analysis of the mouse nidogen-2 gene: nidogen-2 is not essential for basement membrane formation in mice. Mol Cell Biol 22, 6820–6830.

Takagi, Y., Nomizu, M., Gullberg, D., MacKrell, A. J., Keene, D. R., Yamada, Y. and Fessler, J. H. (1996). Conserved neuron promoting activity in Drosophila and vertebrate laminin alpha1. J Biol Chem 271, 18074–18081.

Tanentzapf, G., Devenport, D., Godt, D. and Brown, N. H. (2007). Integrin-dependent anchoring of a stem-cell niche. Nat Cell Biol 9, 1413–1418.

Timpl, R. and Brown, J. C. (1996). Supramolecular assembly of basement membranes. Bioessays 18, 123–132.

Timpl, R., Dziadek, M., Fujiwara, S., Nowack, H. and Wick, G. (1983). Nidogen: a new, self-aggregating basement membrane protein. Eur J Biochem 137, 455–465.

Tsai, P. I., Wang, M., Kao, H. H., Cheng, Y. J., Lin, Y. J., Chen, R. H. and Chien, C. T. (2012). Activity-dependent retrograde laminin A signaling regulates synapse growth at Drosophila neuromuscular junctions. Proc Natl Acad Sci U S A 109, 17699–17704.

Uechi, G., Sun, Z., Schreiber, E. M., Halfter, W. and Balasubramani, M. (2014). Proteomic View of Basement Membranes from Human Retinal Blood Vessels, Inner Limiting Membranes, and Lens Capsules. J Proteome Res.

Urbano, J. M., Torgler, C. N., Molnar, C., Tepass, U., Lopez-Varea, A., Brown, N. H., de Celis, J. F. and Martin-Bermudo, M. D. (2009). Drosophila laminins act as key regulators of basement membrane assembly and morphogenesis. Development 136, 4165–4176.

Wagh, D. A., Rasse, T. M., Asan, E., Hofbauer, A., Schwenkert, I., Durrbeck, H.,Buchner, S., Dabauvalle, M. C., Schmidt, M., Qin, G., et al. (2006). Bruchpilot, a protein with homology to ELKS/CAST, is required for structural integrity and function of synaptic active zones in Drosophila. Neuron 49, 833–844.

Willem, M., Miosge, N., Halfter, W., Smyth, N., Jannetti, I., Burghart, E., Timpl, R. and Mayer, U. (2002). Specific ablation of the nidogen-binding site in the laminin gamma1 chain interferes with kidney and lung development. Development 129, 2711–2722.

Voigt, A., Pflanz, R., Schafer, U. and Jackle, H. (2002). Perlecan participates in proliferation activation of quiescent Drosophila neuroblasts. Dev Dyn 224, 403–412.

Wolfstetter, G. and Holz, A. (2012). The role of LamininB2 (LanB2) during mesoderm differentiation in Drosophila. Cell Mol Life Sci 69, 267–282.

Wolfstetter, G., Shirinian, M., Stute, C., Grabbe, C., Hummel, T., Baumgartner, S., Palmer, R. H. and Holz, A. (2009). Fusion of circular and longitudinal muscles in Drosophila is independent of the endoderm but further visceral muscle differentiation requires a close contact between mesoderm and endoderm. Mech Dev 126, 721–736.

Wu, Z., Sweeney, L. B., Ayoob, J. C., Chak, K., Andreone, B. J., Ohyama, T., Kerr, R., Luo, L., Zlatic, M. and Kolodkin, A. L. (2011). A combinatorial semaphorin code instructs the initial steps of sensory circuit assembly in the Drosophila CNS. Neuron 70, 281–298.

Yao, Y. (2017). Laminin: loss-of-function studies. Cell Mol Life Sci 74, 1095–1115.

Yarnitzky, T. and Volk, T. (1995). Laminin is required for heart, somatic muscles, and gut development in the Drosophila embryo. Dev Biol 169, 609–618.

Yasothornsrikul, S., Davis, W. J., Cramer, G., Kimbrell, D. A. and Dearolf, C. R. (1997). viking: identification and characterization of a second type IV collagen in Drosophila. Gene 198, 17–25.

Yurchenco, P. D. (2011). Basement membranes: cell scaffoldings and signaling platforms. Cold Spring Harb Perspect Biol 3.

Zhu, P., Ma, Z., Guo, L., Zhang, W., Zhang, Q., Zhao, T., Jiang, K., Peng, J. and Chen, J. (2017). Short body length phenotype is compensated by the upregulation of nidogen family members in a deleterious nid1a mutation of zebrafish. J Genet Genomics.

Zhu, X., Ahmad, S. M., Aboukhalil, A., Busser, B. W., Kim, Y., Tansey, T. R., Haimovich, A., Jeffries, N., Bulyk, M. L. and Michelson, A. M. (2012). Differential regulation of mesodermal gene expression by Drosophila cell type-specific Forkhead transcription factors. Development 139, 1457–1466.

